# ITPKB is a conserved regulator of natural killer cell desensitization/education that constrains antitumor immunity

**DOI:** 10.64898/2026.07.12.738073

**Authors:** Yeara Jo, Alexandros Karampatzakis, Seung Won Lee, Alice Peng, Zaina Ghouri, Monsserratt Moya, Elizabeth Andrews, Harrison Sudholz, Chenyu Zhang, Berna Bou Tayeh, Alejandro N. Medina, Jiucheng Ding, Robert A. Saxton, Ziyang Zhang, Yang Shi, Hideho Okada, David H. Raulet

**Affiliations:** Division of Immunology and Molecular Medicine, Department of Molecular and Cell Biology, University of California, Berkeley, Berkeley, CA, USA; Ludwig Institute for Cancer Research, Nuffield Department of Medicine, University of Oxford, Oxford, UK; Department of Chemistry, University of California, Berkeley, Berkeley, CA, USA; Department of Neurological Surgery at University of California, San Francisco, San Francisco, CA, USA; Parker Institute for Cancer Immunotherapy, San Francisco, CA, USA; Helen Diller Family Comprehensive Cancer Center, San Francisco, CA, USA

## Abstract

Natural Killer (NK) cell desensitization induced by persistent stimulation limits durable antitumor immunity, yet the molecular mechanisms governing this dysfunctional state remain poorly defined. To identify conserved regulators of NK cell desensitization, we performed comparative transcriptomic analyses across multiple murine models of persistent activation and dysfunction. This approach defined a shared transcriptional program of desensitization and identified *Itpkb* to be upregulated across multiple distinct contexts. Genetic and pharmacological inhibition of ITPKB enhanced degranulation, cytokine production, and cytotoxicity in both murine and human NK cells under multiple desensitization settings. Mechanistically, ITPKB regulated signaling downstream of persistent activation through the IP_3_/IP_4_ axis, limiting calcium mobilization and NFAT-dependent transcriptional responses in desensitized NK cells. Furthermore, inhibition or deletion of ITPKB enhanced NK-cell mediated tumor control *in vivo* and improved the efficacy of adoptively transferred CAR-NK cells, underscoring the translational potential of targeting this pathway. Together, these findings identify ITPKB as a cell-intrinsic regulator of NK cell desensitization and support targeting the IP_3_/IP_4_ signaling axis to enhance NK cell-mediated antitumor immunity.

## Introduction

Natural killer (NK) cells are innate lymphocytes that play a crucial role in immune surveillance by eliminating transformed or infected cells through direct cytolysis and secretion of proinflammatory cytokines such as IFN-γ and TNF-α^1^. NK cells rely on a repertoire of germline-encoded activating and inhibitory receptors to discriminate healthy from abnormal cells based on molecular cues such as the loss of MHC class I (“missing self”) or expression of stress ligands (“induced self”)^2,3^. Because NK cells mediate rapid cytotoxic responses without requiring substantial clonal expansion and recognize target cells independently of antigen presentation, they have emerged as promising platforms for cancer immunotherapy, particularly in tumors with low mutational burden or poor neoantigen visibility^4^. Multiple approaches have been developed to mobilize endogenous NK cells in tumors, including engineered cytokines^5–8^, activators of innate immunity^9–12^, blockade of inhibitory receptors^13,14^, and bispecific engagers to bridge NK cell activating receptors and tumor-specific molecules^15,16^. Adoptive NK cell therapies, including CAR-NK cells have potential for off-the-shelf applications and are showing promise in the clinic ^10,17–21^. The results suggest that NK cells can reject established tumors if their effector functions can be sustained or enhanced. However, resistance to NK therapy and frequent relapse among responding patients^22,23^ highlight the need for a deeper understanding of the molecular mechanisms that regulate NK cell function in tumors.

NK cell education, sometimes called "licensing", “arming/disarming” or “tuning”, is a quantitative process that tunes NK cell responsiveness depending on their steady state environment^1,24–26^. Interactions of NK cell receptors with MHC and other molecules encountered on surrounding cells establish tolerance to normal ’self’ cells and tunes the capacity of individual NK cells to respond to subsequent stimulation by unhealthy cells, in a manner likened to a rheostat^24–26^. The responsiveness of mature NK cells is only semi-stable, however, and can be decreased or increased when the cells are exposed to greater or lesser steady-state stimulation, respectively^27,28^. This functional plasticity of NK cells is particularly relevant in tumors. For instance, NK cells initially mount strong responses against MHC class I-deficient tumor cells but gradually lose cytotoxic capacity under persistent stimulation, becoming desensitized and unable to control large tumor burdens^5^. Similar desensitization has also been observed under conditions of prolonged engagement with NK-activating ligands^29–31^, suggesting that both “missing self” and “induced self” cues can drive this form of dysfunction. Understanding the molecular mechanisms underlying NK cell desensitization in both steady-state and tumor contexts is therefore critical for designing strategies to reprogram NK cells to sustain antitumor activity.

Analogous to NK cell desensitization, T cells progressively lose effector function upon chronic antigen exposure, entering a state of “exhaustion”^32^. Distinct exhausted T cell states have been defined, including a plastic, progenitor-like population capable of reinvigoration and a more terminally exhausted population that is epigenetically fixed and refractory to rescue^33,34^. Transcriptomic analyses across infection and tumor models have revealed a conserved molecular circuitry driven by chronic stimulation^35,36^, including inhibitory receptors such as PD-1 and LAG-3, blockade of which restores effector activity in the reprogrammable subset of exhausted CD8⁺ T cells^37–41^. Importantly, terminally exhausted T cells exhibit substantial differences in gene expression programs compared to the reprogrammable subset^33,36^. These studies underscore the power of transcriptional profiling to uncover mechanisms of immune dysfunction and identify therapeutic targetable states. In contrast, efforts to define the molecular underpinnings of NK cell desensitization have been more limited, in part due to the diversity of receptor inputs that govern NK cell activation and tolerance. Systematic transcriptomic profiling of NK cells rendered desensitized under distinct conditions may therefore illuminate core regulators of NK cell desensitization and reveal actionable pathways to sustain NK cell antitumor activity.

Based on this rationale, we sought to delineate the transcriptional networks underlying NK cell desensitization using a series of well-controlled, tumor-free models. These reductionist systems recapitulate key features of tumor-associated dysfunction while allowing direct interrogation of intrinsic regulatory mechanisms. Comparative RNA-sequencing of NK cells across multiple desensitization contexts identified both conserved and context-specific transcriptional signatures that collectively define the molecular landscape of NK cell desensitization. Among the recurring differentially expressed genes, *Itpkb*, encoding inositol-trisphosphate 3-kinase B (ITPKB), emerged as a candidate regulator of NK cell desensitization. Although ITPKB has previously been implicated as a negative regulator of NK cell activation and effector function^42^, its role in chronic stimulation-induced NK cell desensitization has not been explored. Here, we combine genetic and pharmacologic perturbation approaches to identify ITPKB as a key mediator of NK cell desensitization arising from persistent stimulation in both murine and human systems. Furthermore, we demonstrate that targeting ITPKB restores NK cell effector function and enhances NK cell-mediated antitumor immunity, including in the context of CAR-NK cell therapy. Together, these findings identify ITPKB as a targetable pathway regulating NK cell desensitization and provide a framework for therapeutically reprogramming dysfunctional NK cells in cancer.

## Results

### Transcriptomic Profiling Identifies Candidate Regulators of NK Cell Desensitization

We set up multiple robust *in vivo* and *in vitro* conditions to compare desensitized NK cells with their functional counterparts. First, we compared functional vs. desensitized NK cells that naturally arise through NK cell education ^24–26^. Normal lymphoid cells lacking MHC I are killed by NK cells, indicating that healthy cells provide stimulatory signals that are normally counterbalanced by inhibitory receptors recognizing self MHC I molecules, such as K^b^ and D^b^ in C57BL6/J (B6) mice. In B6 mice, these receptors include Ly49C and Ly49I (bind to K^b^)^43^, and CD94/NKG2A (binds to a D^b^-derived peptide presented by the Qa-1 nonclassical MHC I molecule)^44^. Individual NK cells can express any combination of Ly49C, Ly49I or CD94/NKG2A (abbreviated C, I and N, respectively), or none of them^26,45,46^. In B6 mice, NK cells lacking all 3 receptors (C^-^I^-^N^-^ NK cells; triple negative, TN) are inhibited the least and therefore persistently stimulated the most by neighboring cells in steady state conditions, resulting in a high degree of desensitization. NK cells expressing only one of the receptors, such as Ly49I^+^ single positive (SP) cells, are only partially inhibited and therefore less desensitized, and triple positive (TP) NK cells are inhibited the most and therefore least desensitized. Consequently, in response to activating receptor engagement *ex vivo*, the degranulation and cytokine production responses follow the pattern TP>SP>TN (Figure 1A, S1A and ^24–26,47,48^). The same pattern held when we gated on the most mature, CD27^-^CD11b+ NK cell subset ^49^ (Figure 1B and S1B). In MHC I-deficient *B2m^−/−^*mice, all NK cells are persistently stimulated and therefore strongly desensitized, similar to TN cells in B6 mice (Figure 1A, S1A, and ^24–26,47^).

**Figure 1.**
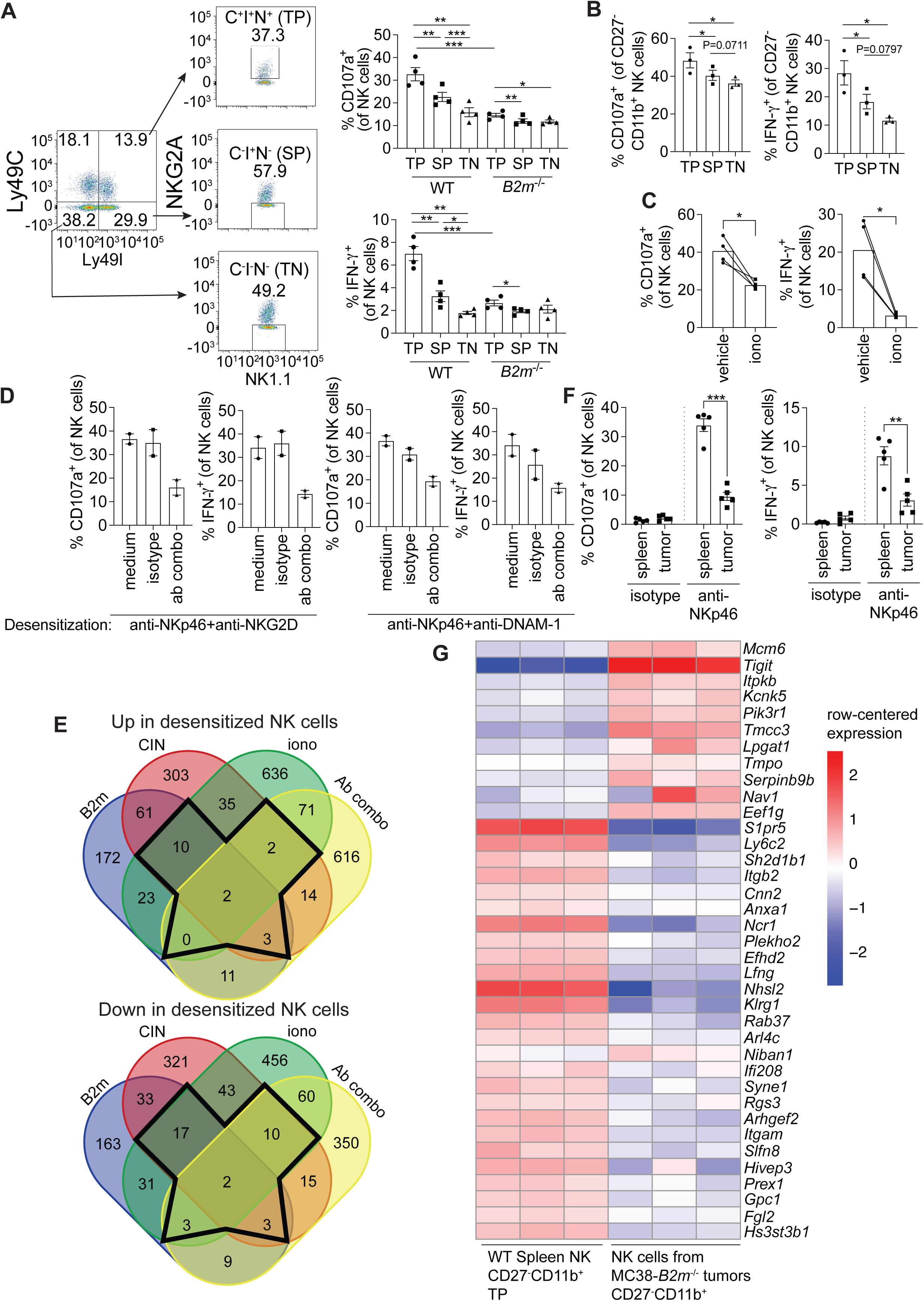
Transcriptomic profiling identifies conserved regulators of NK cell desensitization. (A) Splenocytes from *B2m^+/+^* (WT) and *B2m^−/−^* mice were isolated. Gating strategy for TP (Ly49C⁺Ly49I⁺NKG2A⁺), SP (Ly49C⁻Ly49I⁺NKG2A⁻), and TN (Ly49C⁻Ly49I⁻NKG2A⁻) NK subsets is shown (left). TP, SP, and TN cells from WT and *B2m^−/−^* mice were evaluated for degranulation and IFN-γ production after NKp46 stimulation (right). (B) TP, SP, and TN NK subsets from the CD27⁻CD11b⁺ fraction of WT mice were evaluated for degranulation and IFN-γ production after NKp46 stimulation. (C) WT splenocytes were treated with vehicle or 1 μM ionomycin for 22 hours, then evaluated for degranulation and IFN-γ production following stimulation with NKp46 antibodies. (D) WT splenocytes mice were stimulated for 48 hours with the indicated combinations of plate-bound antibodies or respective isotype controls or left unstimulated and subsequently tested for degranulation and IFN-γ production following NK1.1 stimulation. (E) RNA-seq was performed under multiple conditions: (1) union of differentially expressed genes from three pairwise comparisons among CD27⁻CD11b⁺ NK cells in WT mice: SP vs. TP, TN vs. TP, or TN vs. SP (“CIN”); (2) TP CD27⁻CD11b⁺ NK cells in *B2m^−/−^* vs. WT mice (“B2m”); (3) ionomycin- vs. vehicle-treated NK cells (“iono”); and (4) NK cells stimulated with anti-NKp46 plus anti-NKG2D or anti-DNAM-1 antibodies vs. medium alone (“Ab combo”). Venn diagrams show genes up- (top) or down-regulated (bottom) genes in desensitized vs. functional NK cells. Genes common to ≥3 conditions are outlined in black and listed in Tables 1 and 2. (F) B6 mice were subcutaneously injected with 4 x 10^6^ MC38-*B2m^−/−^* tumor cells and analyzed 10 days later. Splenic and tumor NK cells were stimulated with NKp46 or isotype control antibodies and analyzed for degranulation and IFN-γ production. (G) Heatmaps show genes commonly up- (top) or down-regulated (bottom) in ≥3 desensitization conditions from (E) in the indicated splenic and tumor NK cell populations in normalized rlog-transformed expression data. Rows were centered by subtracting each gene’s mean across samples before plotting; blue indicates lower-than-average expression, white indicates approximately average expression, and red indicates higher-than-average expression. (A-D, F) Bars represent means ± SEM with each symbol representing one mouse. Data are representative of 2-3 independent experiments. Statistical significance was determined using unpaired t-tests (WT TP vs. *B2m^−/−^*TP in (A)) or paired t-tests for the other comparisons. *p < 0.05, **p < 0.01, ***p < 0.001.

To identify conserved molecular pathways associated with NK cell desensitization, we next established additional modes of desensitization, including 22-hour stimulation with ionomycin, a calcium ionophore that mimics receptor-induced Ca^2+^ influx^50^ and is known to induce a state of dysfunction in both T cells and NK cells^51,52^. We confirmed that a 22-hour treatment with ionomycin desensitized murine NK cells (Figure 1C and S1C). As yet another mode of desensitization, we reproduced persistent ‘induced self’ signals by simultaneously stimulating normal NK cells *in vitro* via engagement of multiple activating receptors for an extended period^53^ Thus, NK cells stimulated for 48-hours with plate-bound antibodies targeting NKp46 in combination with NKG2D or DNAM-1, combinations that optimally activate NK cells^54^, were desensitized when subsequently stimulated via a different activating receptor, NK1.1 (Figure 1D and S1D).

Because transcriptional changes are readily measurable and experimentally tractable and bulk RNAseq is more quantitative than single cell RNAseq, we performed bulk RNA-seq to compare desensitized NK cells and their functional counterparts in the aforementioned conditions: 1) SP vs. TP, TN vs. TP, or TN vs. SP NK cells in *B2m*^+/+^ mice (labeled "CIN"), 2) TP NK cells from *B2m*^−/−^ vs. *B2m*^+/+^ mice (B2m), 3) NK cells treated with ionomycin vs. vehicle (iono), and 4) NK cells stimulated for 48 hours with combinations of activating receptor antibodies (Ab combo). Across most conditions, differentially expressed (DE) genes significantly overlapped with literature-reported desensitization-associated transcriptional programs^52,55–57^, supporting the validity of our approach (Figure S1E-H). In addition, gene set enrichment analysis revealed that ionomycin- and persistent activating receptor ligation-induced desensitization shared transcriptional features derived from exhausted CD8^+^ T cells integrated across multiple exhaustion contexts and species^58^, whereas desensitization associated with disruption of MHC class I-mediated inhibitory signaling did not (Figure S1I-L). These findings suggest that distinct desensitization contexts engage partially overlapping molecular programs, with some forms of NK cell desensitization converging at least in part with pathways associated with T cell exhaustion, whereas others likely reflect mechanistically distinct processes.

To find conserved core regulators of NK cell desensitization, we overlapped the DE genes from the 4 comparisons and focused on the genes that were commonly dysregulated in 3 or more comparisons (Figure 1E, black line; Table 1 and 2). These included genes encoding well known regulators of NK cell activity, such as TIGIT, blockade of which augments NK-mediated anti-tumor immunity^13^. However, TIGIT expression alone cannot fully account for NK cell desensitization, as desensitized NK cells show diminished responses even under stimulation conditions that bypass inhibitory interactions between TIGIT and its ligands (CD155 and CD112), such as plate-bound antibody stimulation. This observation suggests the presence of additional cell-intrinsic mechanisms underlying NK cell desensitization.

In MHC I-deficient tumors, NK cells initially recognize the loss of MHC I and attack tumor cells, such that small tumor loads can be eradicated, but with larger tumor loads NK cells eventually become desensitized and lose the capacity to control the tumors^5,59^. Comparing the functional capacity of NK cells in MC38-*B2m*^−/−^ colorectal tumors vs. matched spleens showed significantly compromised degranulation and IFN-γ production by tumor-infiltrating NK cells following NKp46 stimulation (Figure 1F). Gene expression profiling showed that 11 of 17 genes (64.7%) that were commonly upregulated across multiple desensitization conditions were more highly expressed in NK cells from MC38-*B2m*^−/−^ tumors than those from spleens (Figure 1G). We further analyzed the expression of these 17 genes in CD56^dim^CD16^high^ NK cell clusters from a pan-cancer atlas of human tumors^60^. 10 of the 17 genes were more highly expressed in the cluster that exhibited the weakest cytotoxicity ("c6-DNAJB1 cluster") compared to the cluster showing the highest inflammatory score among all NK cell subsets ("c4-NFKBIA cluster") (Figure S1M). Moreover, 26 of 34 genes (76.5%) that were commonly downregulated across multiple desensitization conditions also exhibited reduced expression in desensitized tumor-infiltrating NK cells compared to functional splenic NK cells (Figure 1G).

These findings underscored the strength of our approach in identifying potential core regulators of NK cell desensitization and suggest the utility of our gene set for studying NK cell desensitization in the tumor microenvironment (TME). Together, these results established a foundational gene signature of NK cell desensitization and enabled us to nominate conserved candidate regulators for functional validation in both murine and human systems.

### *Itpkb* promotes functional desensitization in murine NK cells

Among the 17 upregulated genes in the desensitization signature (Figure 1), *Itpkb* was more highly expressed in desensitized NK cells under 3 of the 4 conditions tested: CIN, B2m, and iono (Figure 2A). Additionally, *Itpkb* expression was higher in NK cells infiltrating MC38- *B2m*^−/−^ tumors compared to splenic NK cells (Figure 1G), a finding validated by quantitative(q) RT-PCR (Figure S2A). *Itpkb* encodes inositol-triphosphate 3-kinase B (ITPKB), which phosphorylates soluble inositol(1,4,5)-trisphosphate (IP_3_) to generate inositol(1,3,4,5)– tetrakisphosphate (IP_4_)^61^, thereby modulating signaling downstream of activating receptors. The identification of ITPKB was particularly intriguing because prior studies implicated it as a negative regulator of NK cell effector responses ^42^. We therefore hypothesized that ITPKB upregulation is a mechanism that underlies NK cell desensitization and tested effects of pharmacologic and genetic perturbation of ITPKB in multiple desensitization models.

**Figure 2.**
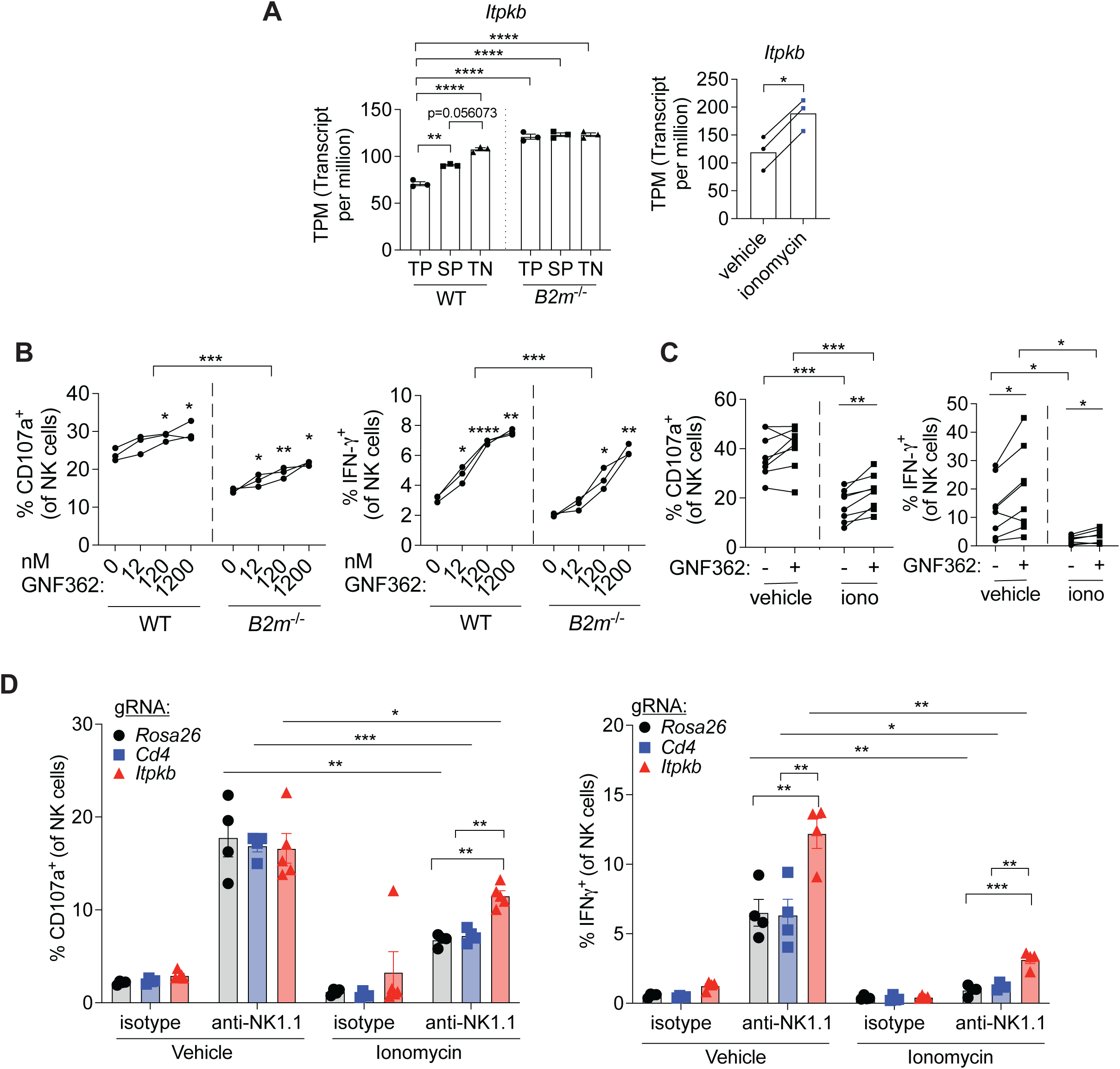
ITPKB promotes functional desensitization in murine NK cells. (A) RNA-seq in Fig. 1E was analyzed for differential expression using DESeq2 with count-based normalization after importing RSEM quantifications through tximport. TPM values for comparisons in approaches (1)-(2) (left) and (3) (right) are shown in plots for visualization only. (B) Splenocytes from WT or *B2m*^−/−^ mice were pre-treated with vehicle or indicated concentrations of ITPKB inhibitor GNF362 for 2 hours, stimulated with NKp46 antibodies in the continued presence of inhibitor and evaluated for degranulation and IFN-γ production. (C) WT splenocytes were treated with vehicle or 1 μM ionomycin for 22 hours in the presence of 50 IU/ml IL-2, pre-treated with vehicle or 1.2 μM ITPKB inhibitor GNF362 for 2 hours, stimulated with NKp46 antibodies in the presence of inhibitor and evaluated for degranulation and IFN-γ production. (D) NK cells electroporated with CRISPR-Cas9 ribonucleoprotein complexes targeting *Itpkb* or, as controls, *Rosa26,* or *Cd4* were treated with vehicle or 1 μM ionomycin for 22 hours, stimulated with NK1.1 antibodies and evaluated for degranulation and IFN-γ production. Error bars represent SEM. Data are representative of 2-3 independent experiments (B, D) or pooled from 2 experiments (C). Statistical significance was determined using Wald test followed by adjustment for multiple testing using the Benjamini-Hochberg method embedded in DESeq2 (A), 2-way ANOVA (WT vs. *B2m^−/−^* in (B)) or paired t-tests for the other comparisons. Each symbol represents an individual mouse. *p<0.05, **p<0.01, ***p<0.001, ****p<0.0001.

To test this while avoiding the developmental defects observed in *Itpkb*-deficient mice, we treated desensitized NK cells and their functional counterparts with a selective and potent ITPKB inhibitor, GNF362^62^, prior to and during functional assays. In desensitized NK cells from *B2m^−/−^* mice, GNF362 treatment resulted in dose-dependent increases in both degranulation and IFN-γ production (Figure 2B). Some enhancement was also observed in NK cells from *B2m^+/+^* mice, possibly reflecting the fact that many NK cells in these mice are partially desensitized^47,48^ or simply reflecting the fact that these NK cells still express some *Itpkb*. Importantly, GNF362 treatment did not compromise cell viability (Figure S2B). GNF362 also enhanced the degranulation responses of WT NK cells that had been desensitized for 22 hours with ionomycin in the presence of low dose IL-2 before restimulating the cells with plate-bound NKp46 antibody (Figure 2C). The inhibitor had no significant effect on NK cells treated with IL-2 alone, perhaps because IL-2 partially alleviated the baseline partial desensitization characteristic of freshly isolated NK cells, Notably, ITPKB inhibition also enhanced IFN-γ production in control NK cells and to a lesser extent in desensitized NK cells (Figure 2C). These findings suggested that ITPKB inhibition enhances the functional activity of ionomycin- desensitized NK cells with a more selective effect on control NK cells. Together, these findings support a role for ITPKB in promoting NK cell desensitization.

In light of the potential off-target effects of pharmacological inhibitors, we deleted *Itpkb* genetically in NK cells using CRISPR (Clustered Regularly Interspaced Short Palindromic Repeats)/Cas9 ribonucleoprotein (RNP) nucleofection. NK cells edited at *Rosa26* (a neutral "safe harbor" locus) or *Cd4* (a gene not expressed in NK cells) were used as controls. Editing efficiency at the *Itpkb* locus was approximately 60%, as determined by Tracking of Indels by Decomposition (TIDE) analysis (Figure S2C)^63^. Genetic deletion of *Itpkb* in NK cells increased degranulation in ionomycin-desensitized NK cells, but not in vehicle-control treated NK cells (Figure 2D, left). In addition, *Itpkb*-edited NK cells exhibited increased IFN-γ production in both control and desensitized conditions (Figure 2D, right) similar to the results with the ITPKB inhibitor.

Notably, *Itpkb*-edited NK cells did not show any significant increases in the levels of granzyme A, granzyme B or perforin compared to *Rosa26*- or *Cd4*-edited control NK cells (Figure S2D). Together, these findings show *Itpkb* deletion enhances granule release and cytokine production by NK cells, rather than increasing effector molecule content, particularly under desensitizing conditions. These results establish ITPKB as a critical mediator of NK cell desensitization and demonstrate that disrupting its activity can partially overcome the loss of effector activity associated with desensitization in mature NK cells.

### ITPKB promotes functional desensitization in a human NK cell line

We next asked whether ITPKB is upregulated in desensitized human NK cells and contributes to functional desensitization. *ITPKB* transcript levels were indeed increased in the NK-92 human NK cell line following ionomycin-induced desensitization (Figure 3A). Furthermore, the ITPKB inhibitor GNF362 enhanced degranulation of both desensitized and control NK-92 cells in response to NKp30 stimulation or co-culture with K562 target cells, in a dose-dependent manner (Figure S3A-B). GNF362 treatments did not induce spontaneous responses in the absence of stimulation (Figure S3C) or affect cell viability (Figure S3D).

**Figure 3.**
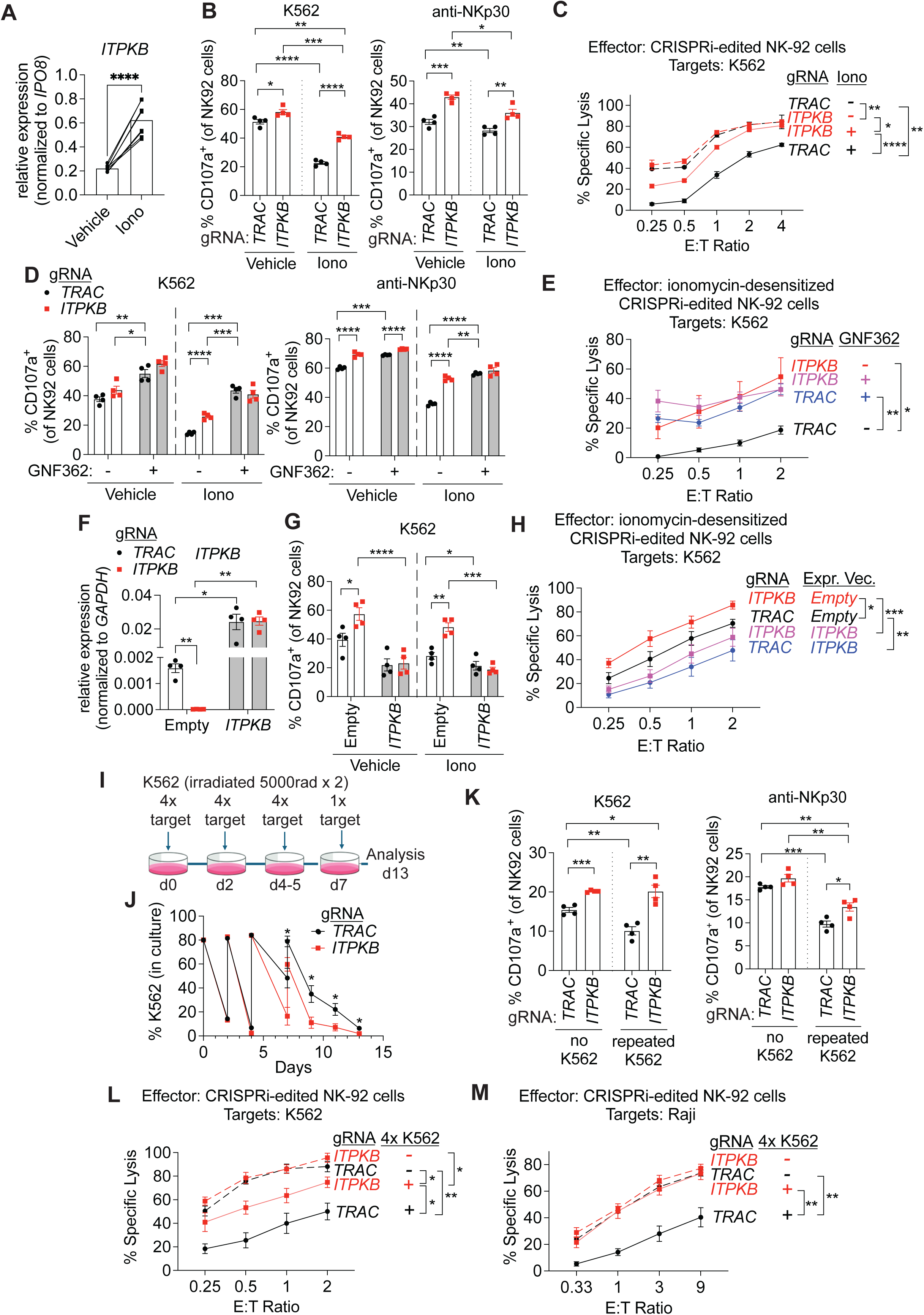
ITPKB promotes functional desensitization in a human NK cell line. NK-92 dCas9-KRAB cells were transduced with lentiviruses encoding dual guide RNAs targeting *ITPKB* or *TRAC* (control), selected with puromycin, and treated with vehicle or 1 μM ionomycin for 22-24 hours before functional assays. (A) *ITPKB* transcript levels were quantified by qRT-PCR in *TRAC*-edited cells. (B-C) Functional responses of control and ionomycin-desensitized cells were assessed by measuring degranulation following stimulation with K562 cells or NKp30 antibodies (B), and by ^51^Cr-release cytotoxicity assays using K562 targets (C). Control cells are shown with dashed lines and desensitized cells with solid lines in (C). (D-E) Cells were pre-treated with vehicle or 1.2 μM ITPKB inhibitor GNF362 for 2 hours and functional assays were performed in the continued presence of inhibitor. Degranulation (D) and cytotoxicity (E) were assessed as in (B-C). (F-H) Cells were additionally transduced with lentiviruses encoding the *ITPKB* open reading frame or an empty vector control. *ITPKB* expression was quantified by qRT-PCR (F), followed by degranulation (G) and cytotoxicity assays (H). (I-M) Cells were desensitized by repeated stimulation with irradiated K562 cells according to the schematic in (I). Remaining K562 cells were quantified over time (J). At 13 days post culture, degranulation following stimulation with K562 cells or NKp30 antibodies (K), and cytotoxicity against K562 (L) or Raji (M) target cells were assessed. Control cells are shown with dashed lines and desensitized cells with solid lines in (L-M). Error bars represent SEM. Data are pooled from 2 (A) or are representative of 2-4 experiments (B-M). Statistical significance was determined using paired t-tests for (A); paired t-tests within *TRAC*- or *ITPKB*-edited cells and unpaired t-tests between *TRAC*- and *ITPKB*-edited cells (B, D, F, G, J and K); or two-way ANOVA for (C, E, H, L, M). Each symbol represents an independently transduced well. *p<0.05, **p<0.01, ***p<0.001, ****p<0.0001.

For a genetic approach, we knocked down *ITPKB* expression using CRISPR interference (CRISPRi) in NK-92 cells stably expressing dCas9-KRAB for transcriptional repression. The cells were transduced with a lentiviral vector encoding two gRNAs targeting *ITPKB*, or two gRNAs targeting *TRAC*, which NK-92 cells don’t express. Compared to *TRAC*- edited cells, *ITPKB*-edited NK-92 cells showed a strong reduction in *ITPKB* expression (Figure S3E) and an increased degranulation in response to K562 cell stimulation in both control and ionomycin-desensitized cells, with a more pronounced enhancement in the desensitized state (Figure 3B). Some impact of *ITPKB* knockdown in nondesensitized cells is not unexpected, because the cells still express *ITPKB*, albeit at lower levels than desensitized cells (Figures 2A and 3A). Degranulation following NKp30 stimulation was also increased under both conditions (Figure 3B). Consistent with enhanced degranulation, *ITPKB*-edited NK-92 cells also exhibited increased cytotoxicity against K562 cells under desensitizing conditions across all effector-to-target ratios tested, with no significant difference observed in control conditions (Figure 3C). Together, these findings indicate that *ITPKB* knockdown improves NK cell degranulation and cytotoxicity after ionomycin-induced desensitization of NK-92 cells.

We next used two complementary approaches to confirm that the enhanced degranulation and cytotoxicity after *ITPKB* knockdown under desensitizing conditions resulted from specific targeting of *ITPKB*, rather than reflecting off-target effects. First, we desensitized both *TRAC*-edited and *ITPKB*-edited NK-92 dCas9-KRAB cells as before and treated them with GNF362 or vehicle alone prior to and during stimulation or during the cytotoxicity assay. GNF362 treatment of *TRAC*-edited controls restored degranulation and cytotoxicity to levels comparable to *ITPKB*-edited cells (Figure 3D-E). Notably, GNF362-treatments resulted in a small but significant increase in degranulation (but not cytotoxicity) even in *ITPKB*-edited cells (Figure 3D), which may reflect inhibition of residual ITPKB from incomplete CRISPRi-mediated knockdown (Figure S3E).

As a second approach to confirm the specificity of CRISPRi-mediated *ITPKB* knockdown, we restored *ITPKB* expression by transducing the cells with a lentiviral expression vector encoding the *ITPKB* open reading frame. The *ITPKB* segment in the vector is expected to be unaffected by the CRISPRi machinery, which targets the *ITPKB* promoter. To validate this construct, we transduced NK-92 cells that do not express the dCas9-KRAB machinery and confirmed effective upregulation of *ITPKB* expression by qRT-PCR (Figure S3F) and reduced degranulation responses (S3G). In *ITPKB*-edited NK-92 dCas9-KRAB cells, *ITPKB* transduction restored high *ITPKB* expression (Figure 3F) and reversed the effects of *ITPKB* knockdown on degranulation and cytotoxicity in both desensitized and vehicle-treated NK cells (Figure 3G-H). These complementary genetic and pharmacologic approaches provide strong evidence that the enhanced degranulation and cytotoxicity observed in desensitized NK-92 cells following *ITPKB* knockdown are due to loss of ITPKB activity, rather than off-target effects of CRISPRi.

Next, we examined the impact of *ITPKB* knockdown on NK cell effector function under a more physiological desensitizing condition. We performed four rounds of repeated challenge with K562 cells, using *TRAC*-edited control and *ITPKB*-edited NK-92 dCas9-KRAB cells^64,65^ (Figure 3I). During the 13-day repeated stimulation period, *TRAC*-edited cells began to lose the capacity to eliminate K562 cells after the third round of challenge on day 5, indicating the onset of desensitization, whereas *ITPKB*-edited cells maintained a greater capacity to eliminate K562 cells (Figure 3J), supporting a role for ITPKB in this form of desensitization. At the end of the 13- day serial co-cultures, the *TRAC*-edited NK cells exhibited reduced degranulation in response to either K562 or NKp30 stimulation compared to cells that were not serially stimulated, whereas *ITPKB*-editing restored much of this difference, especially in K562-stimulated cells (Figure 3K). Consistent with the enhanced degranulation responses, *ITPKB* knockdown also substantially improved cytotoxicity of K562 target cells and Raji target cells by NK-92 cells desensitized with repeated K562 stimulations, but had little or no effect on cytotoxicity by non-desensitized NK-92 cells (Figure 3L-M). Collectively, these complementary pharmacologic and genetic approaches establish ITPKB as a regulator of human NK cell desensitization and demonstrate that its inhibition restores NK cell effector function across multiple desensitization settings.

### ITPKB promotes functional desensitization in primary human NK cells

We next addressed whether ITPKB similarly promotes functional desensitization in primary human NK cells. We isolated NK cells from several peripheral blood donors and expanded them in the presence of irradiated K562 feeder cells expressing membrane-bound IL- 21 and 4-1BB ligand (CSTX002)^66^. Treatments of these NK cells with ionomycin consistently induced desensitization in subsequent responses to stimulation with anti-NKp30, K562 or Raji target cells, and treatments with GNF362 in all cases restored the responses to the levels observed in vehicle-treated control NK cells (Figure 4A), with little or no effect on cell viability (Figure S4A). GNF362 treatments also enhanced the responses of control NK cells, but in most cases to a more limited degree. Notably, GNF362 treatment led to a small but significant increase, or in some cases, a strong trend toward increased IFN-γ production in desensitized NK cells across all stimulation conditions, although levels did not reach those observed in control NK cells (Figure S4B). GNF362 treatments induced little or no response in the absence of stimulation (Figure S4C). Ionomycin also desensitized these NK cells in cytotoxicity assays against Raji target cells (Figure 4B). Killing was partially restored by pharmacological ITPKB inhibition whereas the inhibitor had no significant effect on killing by control NK cells (Figure 4B- D). Together, these findings demonstrate that ITPKB promotes functional desensitization in primary human NK cells and that its inhibition enhances effector functions including degranulation and cytotoxicity.

**Figure 4.**
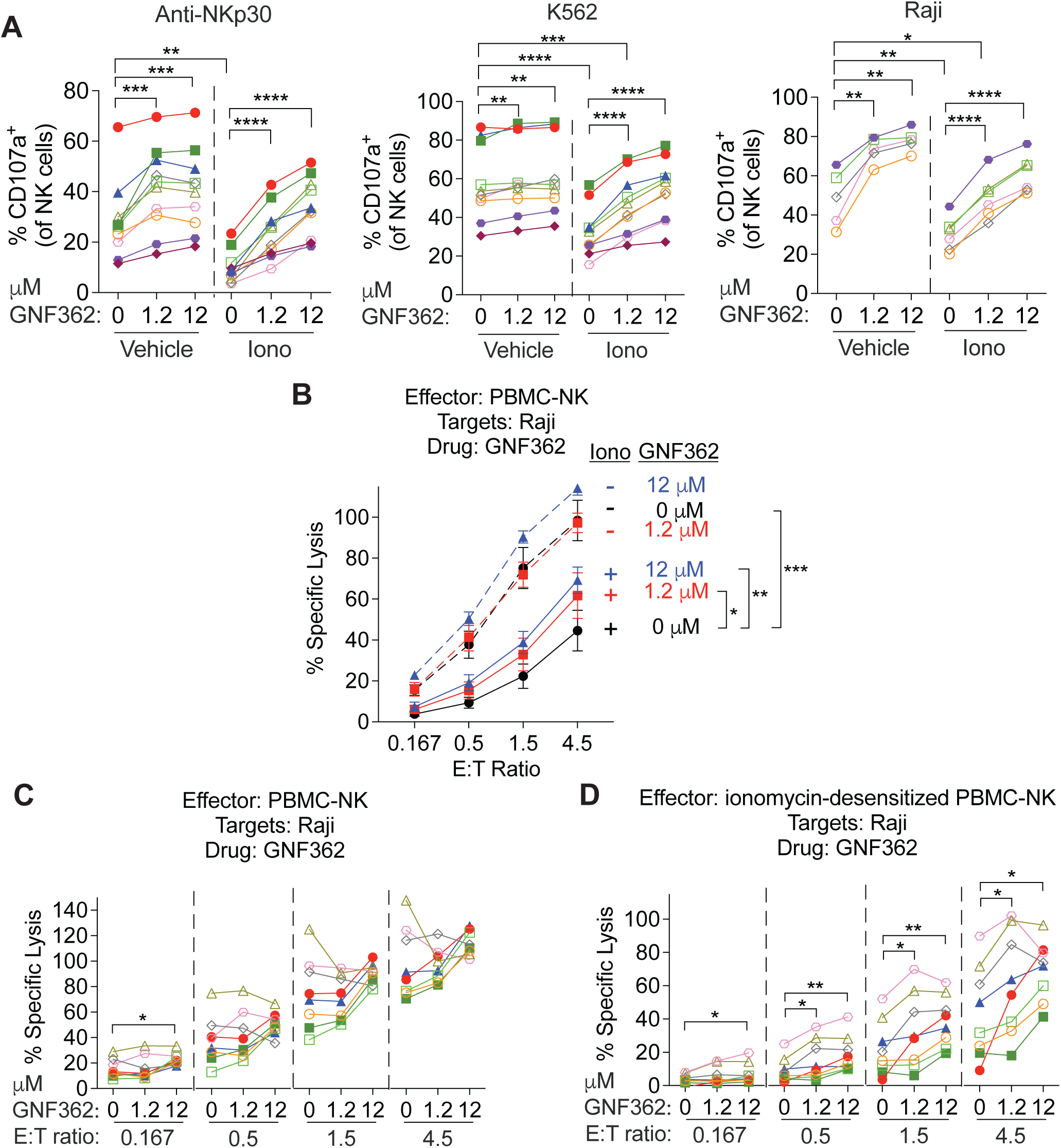
ITPKB promotes functional desensitization in primary human NK cells. Primary NK cells were isolated from human peripheral blood of 10 different donors, indicated with different colors/symbols, and expanded in the presence of irradiated K562 feeder cells expressing membrane-bound IL-21 and 4-1BB ligand (CSTX002). Expanded NK cells were treated with vehicle or 1 μM ionomycin for 16 hours, then pre-treated with vehicle or the indicated concentrations of the ITPKB inhibitor GNF362 for 2 hours. (A) NK cells were stimulated with NKp30 antibodies (left), K562 cells (middle), or Raji cells (right) in the continued presence of GNF362 and analyzed for degranulation. (B-D) Cytotoxicity against Raji cells was assessed with a ^51^Cr-release assay in the continued presence of GNF362. Cytotoxic activities for individual donors from (B) in control (C) or desensitizing (D) conditions are shown. Error bars represent SEM. Data are pooled from 2-3 independent experiments. Statistical significance was determined using paired t-tests (A, C, D) or two-way ANOVA (B). *p<0.05, **p<0.01, ***p<0.001, ****p<0.0001.

### Suppression of IP_4_ generation and enhanced IP_3_R-mediated calcium signaling and NFAT activity restore effector function in desensitized *ITPKB*-deficient NK cells

We sought to define downstream signaling mechanisms through which ITPKB controls degranulation and cytotoxicity in desensitized NK cells. Although the functional effects of ITPKB perturbation extended across multiple desensitization contexts in murine and human NK cells, ionomycin-induced desensitization provides a synchronized and experimentally tractable system for mechanistic interrogation. ITPKB phosphorylates IP_3_ (inositol(1,4,5)-trisphosphate at the 3- position to generate IP_4_ (inositol(1,3,4,5)-tetrakisphosphate)(^61^, Figure S5A), which has been reported to limit NK cell effector activity in murine NK cells^42^. We asked whether it plays a comparable role in mediating ITPKB-dependent desensitization in human NK cells. *TRAC*- edited and *ITPKB*-edited NK-92 dCas9-KRAB cells were desensitized with ionomycin and pretreated with a cell-permeable IP_4_ ester, followed by stimulation with NKp30 antibodies in the continued presence of IP_4_. Supplementation with IP₄ decreased degranulation by *ITPKB*-edited and *TRAC*-edited NK-92 dCas9-KRAB cells under non-desensitizing conditions (Figure 5A). Notably, under desensitizing conditions, a context not examined in previous studies, IP_4_ treatment greatly reduced the enhanced degranulation phenotype conferred by *ITPKB* knockdown, reducing responses to near those of vehicle-treated *TRAC*-edited controls (Figure 5A).

**Figure 5.**
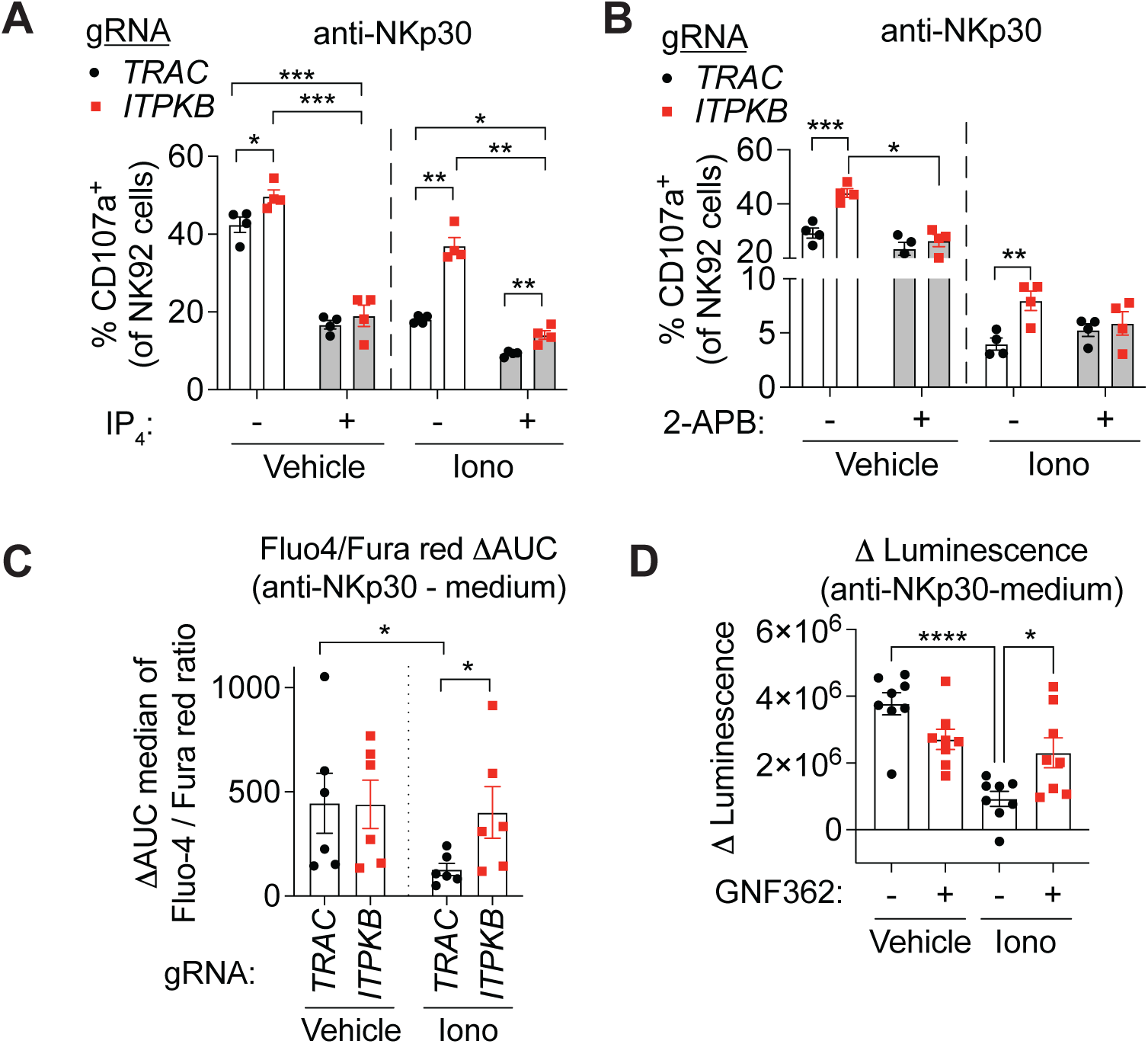
ITPKB-dependent desensitization is associated with accumulation of IP_4_ and decreased IP_3_R-mediated calcium signaling. (A-E) NK-92 dCas9-KRAB cells were transduced with lentiviruses encoding dual guide RNAs targeting *ITPKB* or *TRAC* (control), selected with puromycin, and treated with vehicle or 1 μM ionomycin for 18-24 hours. (A-B) Cells were pre-treated with vehicle or 10 μM cell-permeable IP_4_ (A), or vehicle or 100 μM 2-APB (B), followed by stimulation with NKp30 antibodies in the presence of treatments maintained throughout the assay and analysis of degranulation. (C) Intracellular calcium responses following NKp30 stimulation were measured by Fluo-4/Fura Red ratiometric flow cytometry in vehicle- or ionomycin-treated *TRAC*- or *ITPKB*-edited NK-92 cells. Calcium responses were quantified as the difference in area under the curve (ΔAUC) between anti-NKp30-stimulated and medium control samples. (D) NK-92 cells expressing an NFAT-driven firefly luciferase reporter were treated with vehicle or 1 μM ionomycin for 23 hours, pre-treated with vehicle or 1.2 μM ITPKB inhibitor GNF362 for 2 hours, stimulated with NKp30 antibodies for 9 hours, and analyzed for luciferase activity relative to medium controls. Error bars represent SEM. Data are representative of 2 (A) or 4 (B) independent experiments, or pooled from 4 (C) or 2 (D) independent experiments. Statistical significance was determined using paired t-tests for all comparisons within *TRAC*- or *ITPKB*- edited cells, unpaired t-tests for other cross-group comparisons (A-B), Mann-Whitney test for comparison between *TRAC*- and *ITPKB*-edited cultures (C), or one-way ANOVA with Sidak’s multiple comparisons test (D). Each symbol represents an independently transduced, passaged, and desensitized (A-C) or treated (D) well. *p<0.05, **p<0.01, ***p<0.001, ****p<0.0001.

Additionally, because ITPKB catalyzes the conversion of IP_3_ into IP_4_, it is expected to deplete intracellular IP_3_, which engages IP_3_ receptors (IP_3_Rs) on the endoplasmic reticulum to promote calcium release into the cytosol and activate multiple signaling pathways^67^. Hence, the reduced IP_3_ levels resulting from increased ITPKB action may also contribute to desensitization, whereas enhanced IP_3_R-mediated calcium signaling after depleting *ITPKB* in desensitized NK cells may contribute to the increased effector function. To test this, *TRAC*-edited and *ITPKB*- edited NK-92 dCas9-KRAB cells were desensitized with ionomycin and pretreated with the IP_3_R antagonist 2-APB (2-aminoethyl diphenylborinate), followed by stimulation with NKp30 antibodies in the continued presence of 2-APB. Under desensitizing conditions, 2-APB treatment abolished the enhanced degranulation observed in *ITPKB*-edited cells (Figure 5B). Collectively, these findings suggest that increased IP_4_ generation and decreased IP_3_ availability together act as key downstream mechanisms of desensitization resulting from elevated *ITPKB* expression.

Consistent with this model, alterations in IP_3_/IP_4_ signaling would be expected to affect intracellular calcium dynamics following receptor engagement (Figure S5A). To test this, *TRAC*- or *ITPKB*-edited NK-92 cells were desensitized with ionomycin and loaded with Fluo-4 and Fura Red to track calcium responses following NKp30 stimulation. *TRAC*-edited cells exhibited significantly impaired calcium responses after desensitization compared with vehicle-treated controls (Figure 5C and S5B-C). In contrast, calcium responses were largely preserved following desensitization in *ITPKB*-edited cells and were significantly restored compared to those observed in desensitized *TRAC*-edited cells (Figure 5C and S5B-C). These findings identify impaired receptor-induced calcium signaling as a feature of NK cell desensitization and establish a role for ITPKB in mediating this defect.

Given that calcium signaling is a major determinant of NFAT (nuclear factor of activated T cells) transcription factor activation (Figure S5A), we next assessed whether impaired calcium responses in desensitized NK cells translated into defective NFAT signaling, using NK-92 cells expressing a firefly luciferase reporter under the control of NFAT response elements. Cells were desensitized with ionomycin and pretreated with the ITPKB inhibitor GNF362, followed by stimulation with NKp30 antibodies in the continued presence of GNF362. The stimulation-induced change in NFAT activity was significantly reduced in desensitized NK cells (Figure 5D), largely reflecting an increase in baseline NFAT activity in those cells (Figure S5D-E). GNF362 treatment significantly enhanced NFAT inducibility (Figure 5D), largely restoring the defect observed in desensitized cells. These data suggest that ITPKB promotes NK cell desensitization at least in part by dampening calcium-dependent NFAT activation.

We also examined whether additional signaling pathways downstream of NK-activating receptors were similarly affected. Phosphorylation of Akt/protein kinase B, previously reported to be suppressed by IP_4_ in murine NK cells under non-desensitized conditions^42^, did not differ significantly between desensitized and non-desensitized NK cells and was not further enhanced by *ITPKB* knockdown (Figure S5F). In contrast, ERK phosphorylation following NKp30 stimulation was significantly reduced in desensitized NK cells (Figure S5G-H), consistent with prior observations in tumor-infiltrating NK cells^5^. However, it was not significantly restored by *ITPKB* knockdown (Figure S5G-H), in agreement with previous findings in murine *Itpkb*^-/-^ NK cells^42^. Collectively, these results suggest that ITPKB promotes NK cell desensitization primarily by limiting IP_3_-dependent calcium signaling rather than by broadly suppressing downstream signaling pathways.

### Deletion or inhibition of ITPKB enhances NK cell-mediated tumor control *in vivo*

The increased *Itpkb* transcript levels in tumor-infiltrating mouse NK cells (Figure 1G and S2A), likely applies to NK cells in at least some human tumors. Published single-cell RNA-seq datasets using the scDVA database^60^ showed enrichment of *ITPKB* expression in NK cells in tumors compared to matched normal tissues in colorectal cancer, breast cancer, hepatocellular carcinoma, and pancreatic cancer, whereas some other tumor types exhibited comparable or lower expression relative to normal tissues (Figure S6A). These findings suggested that elevated ITPKB may contribute to NK cell desensitization in several types of human cancer.

To determine whether extinguishing *Itpkb* expression could enhance NK cell-mediated tumor control *in vivo*, we disrupted *Itpkb* in murine NK cells using CRISPR/Cas9 RNP nucleofection as described in Figure 2, and again achieved approximately 60% knockout of the *Itpkb* gene. Edited NK cells were adoptively transferred to *Rag2^−/−^Il2rg^−/−^* recipient mice, which lack T, B, and NK cells followed by a single injection of H9-MSA, a half-life extended IL-2 ’superkine’ with enhanced affinity for IL-2Rβ/γ complexes^10,68^. Four days after NK cell transfer, recipient mice were implanted subcutaneously (s.c.) with MHC I-deficient MC38-*B2m*^−/−^ colorectal cancer cells. *Itpkb*-editing of NK cells significantly improved tumor control compared to editing of *Rosa26* (Figure S6B-C). The effect was reproducible in independent experiments but modest, likely reflecting incomplete CRISPR editing and the fact that the NK cells were not previously desensitized.

To address the role of ITPKB in a scenario where NK cells were previously desensitized, we tested pharmacologic inhibition of ITPKB on NK-mediated control of established MC38-*B2m*^−/−^ tumors at a time point (day 10 after tumor implantation) when intratumoral NK cells were demonstrably desensitized (Figure 1F). Tumors were initiated in *Rag2^−/−^* mice to exclude effects of T and B cells, and the mice were administered GNF362 or vehicle by oral gavage on days 9, 12 and 14. ITPKB inhibition significantly delayed tumor growth compared to vehicle-treated controls (Figure 6A and S6D). Notably, antibody-mediated depletion of NK cells abrogated the difference in tumor growth between GNF362- and vehicle-treated mice, confirming that the enhanced tumor control was NK cell-dependent (Figure 6A and S6D). Furthermore, NK cells in day 13 tumor dissociates from GNF362-treated mice, when tumor sizes were still comparable to control tumors, showed elevated degranulation and cytokine responses compared to NK cells in vehicle-treated tumors after *ex vivo* NK1.1 stimulation but not without *ex vivo* stimulation (Figure 6B). Elevated responses were also observed in splenic NK cells from GNF362-treated mice (Figure S6E). GNF362 treatment did not significantly alter the frequency or staining intensity of granzyme A-, granzyme B-, or perforin-expressing NK cells (Figure S6F-H). Together, these findings demonstrated that pharmacological inhibition of ITPKB enhances NK cell-mediated tumor control by restoring granule release and cytokine production in desensitized NK cells within the TME, rather than by increasing effector molecule content.

**Figure 6.**
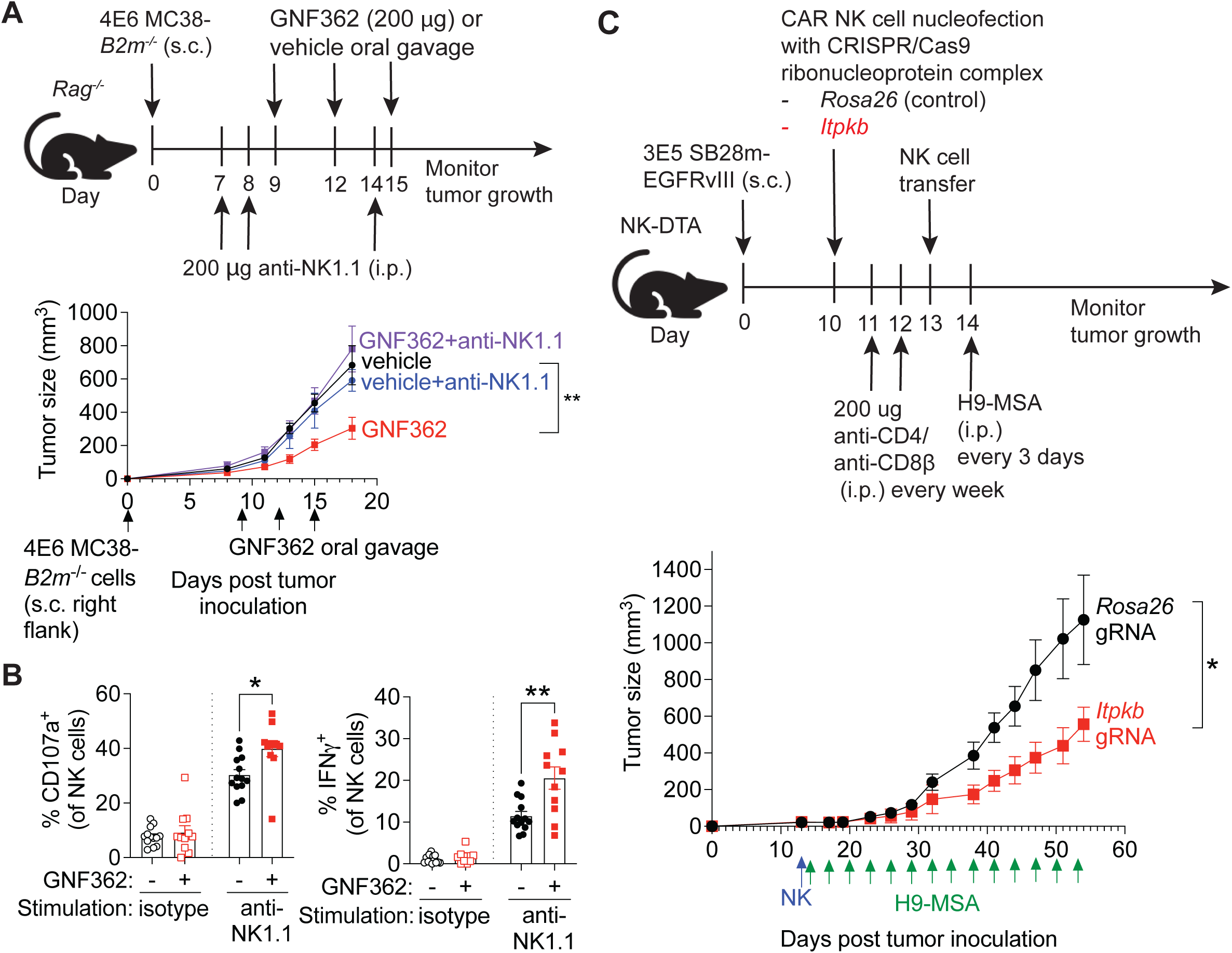
Deletion or inhibition of *ITPKB* enhances NK cell-mediated tumor control *in vivo*. (A) Experimental schematic and tumor growth kinetics following treatments of established tumor-bearing mice with ITPKB inhibitor GNF362. (B) Tumor-infiltrating lymphocytes isolated on day 13 from mice treated as in (A) were stimulated ex vivo with isotype control or NK1.1 antibodies in the presence (or absence) of GNF362 and analyzed for degranulation and IFN-γ production in gated NK cells. (C) Experimental schematic and tumor growth kinetics assessing the impact of *Itpkb* deletion on CAR-NK cell-mediated control of subcutaneous SB28m-EGFRvIII glioma tumors. Error bars represent SEM. Data are representative of two independent experiments (A), pooled from two independent experiments (B), or from one significant experiment with a second experiment showing a p value of 0.086 (C). Statistical significance was determined by two-way ANOVA (A, C) or unpaired t-tests (B). Each symbol represents an individual mouse. *p < 0.05, **p < 0.01.

Given that both genetic deletion and pharmacological inhibition of ITPKB enhanced NK cell effector function, we next examined whether we could harness this property to generate more effective NK cells for cellular therapies, such as chimeric antigen receptor (CAR)-NK cells. We s.c. inoculated syngeneic SB28 glioblastoma cells expressing epidermal growth factor receptor variant III (EGFRvIII)^69^ into NK-DTA mice, which specifically lack NK cells^70^. C57BL/6J- background transgenic mice carrying a murinized anti-EGFRvIII CAR downstream of a Lox-Stop-Lox cassette inserted in the *Rosa26* locus^69^ were crossed with *Vav-Cre* mice to activate CAR expression in hematopoietic cells. CAR-expressing NK cells were expanded from splenocytes of these mice, purified, and nucleofected with Cas9 RNPs containing either *Itpkb*- or *Rosa26*-targeting gRNA and adoptively transferred into tumor-bearing mice on day 13 post tumor inoculation. To ensure that any observed antitumor effects were mediated by the transferred NK cells, recipient mice were depleted of CD4⁺ and CD8⁺ T cells. Starting one day after NK cell transfer, all recipient mice received repeated doses of H9-MSA to support NK cell persistence and proliferation. In this setting, *Itpkb*-edited CAR NK cells conferred a substantial improvement in tumor control compared to *Rosa26*-edited CAR NK cells (Figure 6C and S6I), supporting the potential of engineering ITPKB-deficient NK cells as a therapeutic strategy to achieve sustained antitumor activity.

Together, these findings demonstrate that genetic deletion or pharmacological inhibition of ITPKB enhances NK cell-mediated tumor control by restoring effector function under desensitized conditions and sustaining antitumor activity, providing evidence that *ITPKB* upregulation represents a significant mechanism of NK cell desensitization, and highlighting ITPKB as a promising target for NK cell-based immunotherapy.

## Discussion

Cancer remains the second leading cause of death in the United States^71^. NK cells possess potent antitumor effector functions and hold great promise for cancer immunotherapy^1^. However, NK cells frequently exhibit functional impairment within tumors, which correlates with adverse clinical outcomes^72,73^. Chronic stimulation within the TME progressively desensitizes NK cells and impairs their capacity to control tumor growth. Yet, the molecular basis of NK cell desensitization remains poorly understood and represents one of the most important unresolved questions in NK cell biology. Here, through comparative transcriptomic profiling across multiple tumor-free models of NK cell desensitization, we identified a set of conserved genes dysregulated during desensitization. Among these candidates, genetic and pharmacological suppression of ITPKB enhanced NK cell effector function and improved NK cell-mediated tumor control, particularly under desensitized conditions in murine and human NK cells. Mechanistically, ITPKB restrained NK cell activation through the IP_3_/IP_4_ axis, limiting calcium flux. These findings establish ITPKB as a molecular brake that enforces NK cell desensitization and reveal a targetable pathway for enhancing NK cell-mediated antitumor immunity.

Our transcriptomic profiling of across multiple tumor-free desensitization models provides a framework for identifying core regulators of NK cell dysfunction while minimizing confounding NK cell-extrinsic influences. Previous studies of NK cell education and desensitization have emphasized non-transcriptional mechanisms, including lysosomal remodeling^74^, spatial receptor organization ^55^, and phosphatase localization at the immunological synapse^56^, whereas studies examining transcriptional regulators have largely focused on single experimental contexts, such as individual tumor models^75^, chronic receptor stimulation^57^, or cytokine-driven suppression^76^. Furthermore, transcriptomic analyses of tumor-infiltrating NK cells are often confounded by metabolic stress, hypoxia, stromal interactions, and immunosuppressive cytokines within the TME^77^, making it difficult to distinguish cell-intrinsic desensitization programs from broader environmental effects. By integrating multiple reductionist models of persistent stimulation, we identified conserved dysregulated genes associated with NK cell desensitization. Notably, several genes identified through this approach, including *Tigit, S1pr5, and Ly6c2*, were also dysregulated in NK cells from an independent model of chronic myeloid leukemia^75^, supporting the broader relevance of the transcriptional programs identified here. Importantly, this approach enabled efficient perturbation-based validation and identified ITPKB as a conserved regulator of NK cell desensitization with translational relevance to anti-tumor immunity.

While our analyses identified a core set of genes recurrently dysregulated across desensitization models, many transcriptional changes were context-specific, suggesting that NK cell desensitization arises through overlapping but distinct molecular programs. Consistent with this concept, transcriptional signatures associated with exhausted CD8^+^ T cells were enriched in NK cells desensitized by ionomycin treatment or prolonged stimulation with combinations of activating antibodies, but not in desensitization induced by altered inhibitory receptor signaling. Prior studies in the NK cell education field similarly identified elevated expression of *Ptpn6*, encoding the phosphatase SHP-1, in NK cells desensitized by chronic stimulation resulting from reduced inhibitory receptor engagement, establishing SHP-1 as a positive regulator of NK cell desensitization^56,78^. Furthermore, we found that impaired ERK activation remained largely resistant to ITPKB modulation. Together, these findings support the concept that multiple mechanisms contribute to NK cell desensitization. Importantly, our comparative transcriptomic dataset provides a resource for systematically identifying both conserved and context-specific regulators of NK cell dysfunction beyond ITPKB. It will also be important to determine whether the candidate regulators identified through this framework contribute to NK cell dysfunction induced by other suppressive conditions, including chronic exposure to IL-15^79^ or immunosuppressive cytokines such as TGF-β^80^.

Our data indicate that *Itpkb* is upregulated under conditions of strong and chronic activation, such as ionomycin treatment or chronic stimulation in Ly49C^-^Ly49I^-^NKG2A^-^ (TN) and *B2m^−/−^* NK cells, but not following prolonged stimulation with combinations of activating antibodies. Given that ionomycin directly elevates intracellular Ca^2+^ and TN and *B2m^−/−^* NK cells experience chronic stimulation that involves both “missing self” and “induced self” recognition, *Itpkb* induction may depend on a signaling context generated by both persistent activation and diminished inhibitory signaling or may require a longer duration of stimulation than achieved in this system. This raises the possibility that *Itpkb* expression is part of a feedback circuit initiated by chronic calcium signaling, acting to restrain further activation and limit excessive calcium signaling. Prior studies demonstrating elevated Ca^2+^ and impaired survival in ITPKB-deficient T and B cells^62,81,82^ further support this possibility. Importantly, however, pharmacological inhibition or genetic deletion of ITPKB in our systems was well tolerated and enhanced NK cell effector function and antitumor responses with little or no associated cell death. Together, these findings propose a model in which chronic calcium signaling induces ITPKB expression as an adaptive negative feedback mechanism that restrains NK cell activation and contributes to NK cell desensitization.

Our findings further suggest that NK cell desensitization exists along a spectrum of partially reversible dysfunctional states. Previous studies have demonstrated substantial functional plasticity of mature NK cells, as NK cells from *B2m^−/−^* donors transferred into MHC I-expressing WT hosts regain responsiveness^27,28^. Similarly, cytokine-mediated reinvigoration of desensitized tumor-infiltrating NK cells has been shown to temporarily restore effector function and confer therapeutic benefit^5,9,10^. Consistent with this concept, pharmacologic inhibition of ITPKB restored effector function even in already desensitized NK cells across multiple experimental contexts, supporting the reversibility of at least some desensitized states. Although the effects of ITPKB modulation were most pronounced in desensitized NK cells, more modest enhancements were observed in some non-desensitized murine and human NK cells consistent with the fact that *ITPKB* expression was reduced but still present in those non-desensitized cells. In experimental settings involving exposure to activating cytokines like IL-2, such as our CRISPR-mediated editing studies using Cas9 RNP nucleofection, ITPKB modulation had more limited effects on baseline NK cell function, possibly reflecting alleviated desensitization and/or reduced ITPKB expression or activity. Together, these findings suggest that NK cell desensitization encompasses variable states that differ in their dependence on ITPKB-mediated regulation and capacity for functional recovery. However, genome-wide changes in DNA methylation have been reported in dysfunctional NK cells ^83^. It is plausible that, analogous to exhausted T cells, where populations defined by fixed epigenetic states are refractory to reinvigoration^34,84^, a similarly stable, less reversible desensitized state exists in NK cells, as we have proposed elsewhere^85^. This raises the possibility that more terminal or epigenetically stabilized desensitized states exist in which ITPKB inhibition alone is insufficient to fully restore NK cell function.

Our findings also have important translational implications for cancer immunotherapies. While systematic inhibition of ITPKB enhanced NK cell-mediated tumor control in our models, the broader immunologic consequences of therapeutic ITPKB inhibition will require careful consideration. Notably, ITPKB confers resistance to various chemotherapies in multiple tumor models by directly acting within tumor cells^86,87^. Thus, systemic ITPKB inhibition could enhance tumor control by simultaneously augmenting NK cell activity and sensitizing tumor cells to cytotoxic therapies. However, prior studies demonstrated that ITPKB inhibition can cause activation-induced cell death of T cells and impair development of T and B cells ^62,81,82^. Our *in vivo* studies with the inhibitor were therefore performed in *Rag2^−/−^* mice which lack T and B cells to avoid this potential problem. In addition, ITPKB has been implicated in regulating activation^88^ and CCR7 expression in dendritic cells^89^ as well as longevity and function of hematopoietic stem cells^90^, raising the possibility that sustained systemic inhibition could produce context-dependent immunologic or hematologic toxicities. These observations suggest that strategies enabling selective or transient modulation of ITPKB in NK cells may provide a feasible therapeutic approach to enhance antitumor immunity while minimizing broader toxicities.

In summary, these findings identify ITPKB as a key regulator that imparts and sustains NK cell desensitization and support efforts to target ITPKB and other negative feedback programs in NK cells for therapeutic applications.

## Supplementary Figure Legends

**Supplementary Figure 1. Comparison of NK cell desensitization gene signatures with previously reported datasets, related to Figure 1**.

(A-D) Isotype control stimulation conditions corresponding to Figure 1A-D are shown.

(E) RNA-seq analysis comparing TP (Ly49C⁺Ly49I⁺NKG2A⁺) CD27⁻CD11b⁺ NK cells in *B2m^−/−^*vs. WT mice. Venn diagrams show the overlap of significantly down- or up-regulated genes identified in this study with those identified in a published microarray dataset^55^.

(F) RNA-seq analysis comparing SP (Ly49C⁻Ly49I⁺NKG2A⁻) vs. TP, TN (Ly49C⁻Ly49I⁻NKG2A⁻) vs. TP, or TN vs. SP CD27⁻CD11b⁺ NK cells in WT mice. Differentially expressed genes (DEGs) were defined as the union of DEGs from the three pairwise comparisons. Venn diagrams show overlap of significantly down- or up-regulated genes between this study and a published RNA-seq dataset^56^.

(G) RNA-seq analysis comparing NK cells treated with 1 μM ionomycin or vehicle for 22 hours. Venn diagrams show overlap of down- or up-regulated genes between this study and a published microarray dataset^52^.

(H) RNA-seq analysis comparing NK cells stimulated for 48 hours with plate-bound anti-NKp46 in combination with anti-NKG2D or anti-DNAM1, vs. cells maintained in medium alone. DEGs were defined as the union of DEGs from the two antibody-combination vs. medium comparisons. Venn diagrams show overlap of down- or up-regulated genes between this study and a published RNA-seq dataset^57^.

(I-L) Gene set enrichment analysis using T cell exhaustion signatures from Zhang, et al. 2022^58^ was applied to DEGs derived from ionomycin-treated versus vehicle-treated NK cells in panel G (I), anti-NKp46 + anti-NKG2D-stimulated vs. medium-treated NK cells in panel H (J), panel E (K), and the TN vs. TP comparison in panel F (L).

(M) For the genes upregulated in ≥3 desensitization conditions, the geometric means of their expression levels in each cell of the published scRNAseq data as reported in the scDVA database (http://pan-nk.cancer-pku.cn/) were calculated and used to compute a signature score as shown in the boxplot (left); higher values indicate higher combined expression of the signature. Cluster median expression for each gene was z-scored across groups (right). Heatmap colors show z-scored cluster median expression for each gene across groups (red = z-score of 1, higher relative expression; blue = z-score of -1, lower relative expression).

(A-D) Bars represent means ± SEM with each symbol representing one mouse. Data are representative of 2-3 independent experiments. Statistical significance was determined using unpaired t-tests (WT TP vs. *B2m^−/−^* TP in (A)), Fisher’s exact tests (E-H), or paired t-tests for the other comparisons. *p < 0.05, **p < 0.01.

**Supplementary Figure 2. Comparison of *Itpkb* expression in NK cells from spleen versus tumors and assessment of CRISPR editing efficiency, related to Figure 2**.

(A) C57BL/6J mice were subcutaneously injected with 4 x 10^6^ MC38-*B2m^−/−^*tumor cells and analyzed 18 days post inoculation. NK cells were FACS-purified from spleens and tumors and *Itpkb* expression was quantified by qPCR. Expression levels were normalized to *Rpl13a* (left) or *Tbp* (right).

(B) Splenocytes from WT or *B2m*^−/−^ mice were pre-treated with vehicle or indicated concentrations of ITPKB inhibitor GNF362 for 2 hours, stimulated with NKp46 antibodies in the presence of inhibitor, and assessed for cell viability.

(C-D) NK cells were electroporated with CRISPR-Cas9 ribonucleoprotein (RNP) complexes containing *Itpkb-* or *Rosa26*-targeting gRNAs and cultured for 3 days. Genome editing efficiency was assessed using Tracking of Indels by Decomposition (TIDE) analysis^63^ (C). Gated NK cells treated with vehicle or 1 μM ionomycin for 22 hours were assessed for granzyme A, granzyme B, and perforin proteins by flow cytometry.

Error bars represent SEM. Data are pooled from 2 independent experiments (A, C) or representative of 2 independent experiments (B, D). Statistical significance was determined using paired t-tests and Wilcoxon signed rank tests (A), paired t-tests for comparisons within WT or *B2m^−/−^* cells in (B) and (D), or 2-way ANOVA for the comparison of WT vs. *B2m^−/−^* in (B). Each symbol represents an individual mouse. *p<0.05.

**Supplementary Figure 3. Effects of pharmacological inhibition, CRISPRi-mediated knockdown, and overexpression of *ITPKB* on *ITPKB* expression and degranulation n a human NK cell line, related to Figure 3**.

(A-D) NK-92 cells were treated with vehicle or 1 μM ionomycin for 23 hours and pre-treated with vehicle or the indicated concentrations of ITPKB inhibitor GNF362 for 2 hours. Cells were stimulated with NKp30 antibodies (A) or K562 cells (B), or cultured in medium alone (C) in the continued presence of inhibitor, and analyzed for degranulation. Cell viability under the same stimulation conditions as in (A) and (B) was assessed in (D).

(E) NK92 dCas9-KRAB cells were transduced with lentiviruses encoding dual guide RNAs targeting *ITPKB* or *TRAC* (negative control), followed by puromycin selection. *ITPKB* transcript levels were quantified by qRT-PCR.

(F-G) NK-92 cells were transduced with lentiviruses encoding the *ITPKB* open reading frame or an empty vector control. *ITPKB* transcript levels were quantified by qRT-PCR (F) and the degranulation were determined after stimulation with NKp30 antibodies or K562 target cells (G). Error bars represent SEM. Data are representative of 2-4 experiments. Statistical significance was determined using paired t-tests for (A-D) or unpaired t-tests for (E-G). Each symbol represents an independently transduced well. *p<0.05, **p<0.01, ***p<0.001.

**Supplementary Figure 4. Pharmacological inhibition of ITPKB enhances effector cytokine production and cytotoxic activity in control and desensitized primary human NK cells, with minimal effects on spontaneous activation or viability, related to Figure 4**.

Primary NK cells were isolated from human peripheral blood and expanded in the presence of irradiated K562 feeder cells expressing membrane-bound IL-21 and 4-1BB ligand (CSTX002). Expanded NK cells were treated with vehicle or 1 μM ionomycin for 16 hours, then pre-treated with vehicle or the indicated concentrations of the ITPKB inhibitor GNF362 for 2 hours. NK cells were stimulated with NKp30 antibodies (left), K562 cells (middle), or Raji cells (right) and analyzed for viability (A) and IFN-γ production (B). NK cells cultured in medium alone were analyzed for spontaneous degranulation (left) and IFN-γ (right) production (C). Error bars represent SEM. Data are pooled from 2-3 independent experiments. Statistical significance was determined using paired t-tests where each symbol represents an independent blood donor. *p<0.05, **p<0.01, ***p<0.001, ****p<0.0001.

**Supplementary Figure 5. Desensitized NK cells exhibit elevated baseline NFAT activity and impaired ERK activation upon stimulation independent of ITPKB perturbation, related to Figure 5**.

(A) Proposed mechanism by which ITPKB promotes NK cell desensitization.

(B-C) Representative intracellular calcium flux measured as the Fluo-4/Fura Red fluorescence ratio in TRAC- or ITPKB-edited NK-92 cells treated with vehicle (B; control) or ionomycin (C) to induce desensitization. Cells were stimulated with anti-NKp30 antibody (blue) or maintained in medium alone (red), and intracellular calcium responses were monitored by flow cytometry. Traces represent the median Fluo-4/Fura Red ratio over time.

(D-E) NK-92 cells expressing an NFAT-driven firefly luciferase reporter were treated with vehicle or 1 μM ionomycin for 23 hours to induce desensitization, followed by pre-treatment with vehicle or 1.2 μM ITPKB inhibitor GNF362 for 2 hours. Cells were then left unstimulated (medium control; D) or stimulated with NKp30 antibodies (E) for 9 hours. Luciferase activity was measured.

(F-H) NK92 dCas9-KRAB cells transduced with lentiviruses encoding dual guide RNAs targeting *ITPKB* or *TRAC* (control) were treated with vehicle or 1 μM ionomycin for 22-24 hours, rested for 2 hours, and stimulated with NKp30 antibodies for 10 (F) or 30 (G-H) minutes. Akt (F) and ERK phosphorylation (G-H) were assessed by flow cytometry and fold change relative to unstimulated baseline was quantified. Representative histograms of *TRAC*-edited (black) or *ITPKB*-edited (red) cells for ERK phosphorylation are shown in (G), with control cells indicated by solid lines and desensitized cells by shaded histograms. Error bars represent SEM. Data are pooled from 2-3 independent experiments. Each symbol represents an independently treated sample (D, E) or an independently transduced well (F, H). **p<0.01, ***p<0.001.

**Supplementary Figure 6. ITPKB inhibition enhances NK cell function in the spleen but does not alter granzyme A/B or perforin expression in tumor-infiltrating NK cells, related to Figure 6**.

(A) Published single cell RNA-seq datasets were analyzed using the scDVA database to compare *ITPKB* expression in NK cells from tumors and matched normal tissues across indicated cancer types. Violin plots depict normalized *ITPKB* expression in NK cells from normal tissues (red) and tumors (blue).

(B) Experimental schematic and tumor growth kinetics following adoptive transfer of purified NK cells that were edited for *Itpkb* or *Rosa26*.

(C-D) Spider plots corresponding to (B) and Figure 6A, respectively.

(E) Splenic lymphocytes from the experiment in Figure 6A (without NK-depletion) were isolated on day 13 after tumor inoculation, pretreated with vehicle or 1.2 µM GNF362 for 1 hour, stimulated with isotype control or NK1.1 antibodies for 5 hours in the continued presence of GNF362, and analyzed for degranulation (left) and IFN-γ production (right) by NK cells.

(F-H) Tumor-infiltrating lymphocytes from the same experiment were isolated on day 13 after tumor inoculation, pretreated with vehicle or 1.2 µM GNF362 for 1 hour, and cultured in medium for 5 hours in the continued presence of GNF362 without additional stimulation. Frequencies of granzyme A⁺, granzyme B⁺, and perforin⁺ (left) NK cells and the corresponding geometric mean fluorescence intensities (right) were quantified by flow cytometry.

(I) Spider plots corresponding to Figure 6C. Error bars represent SEM. Data are representative of two independent experiments (B, F-H) or pooled from two experiments (E). Statistical significance was determined by two-way ANOVA (B) or unpaired t-tests (E-H). Each symbol represents one mouse. *p < 0.05; **p < 0.01.

## Materials and Methods

### Mouse strains

Mice were maintained under specific pathogen–free conditions at the University of California (UC) Berkeley. C57BL/6J (B6), *Rag2*^−/−^, and *B2m*^−/−^ mice were purchased from the Jackson Laboratory. *Rag2*^−/−^ *Il2rg*^−/−^ mice were purchased from Taconic Biosciences. *Ncr1^iCre^* mice on the B6 background were generously provided by E. Vivier. NK-DTA mice were generated by breeding *Ncr1^iCre^*mice to B6-*Rosa26^LSL-DTA^* mice (Jackson Laboratory). Transgenic mice carrying a knock-in chimeric antigen receptor recognizing epidermal growth receptor variant III (CAR KI mice) were provided by H. Okada. CAR KI mice were bred with the *Vav^iCre^* mice (JAX 008610) to drive hematopoietic expression of the CAR construct. Offspring were genotyped by PCR for expression of both transgenes. For all breeding and experimental cohorts, mice were maintained hemizygous for Cre to minimize any potential effects of the Cre random insertion. Both male and female mice 8-24 weeks of age were used unless otherwise indicated.

All animal studies were conducted in accordance with protocols approved by the UC Berkeley Institutional Animal Care and Use Committee (IACUC).

### Cell lines and culture conditions

MC-38 *B2m*^−/−^ cells (obtained from Dragonfly Therapeutics), SB28-mEGFRvIII cells (provided by H. Okada), and HEK293T cells (UC Berkeley Cell Culture Facility) were cultured in Dulbecco’s modified Eagle’s medium (DMEM; Thermo Fisher Scientific). NK-92 cells (American Type Culture Collection; ATCC; male origin), NK-92 dCas9-KRAB cells (Laboratory for Genomics Research, University of California, San Francisco; male origin), and NK-92 cells expressing luciferase reporter for NFAT activity (provided by Z. Zhang; male origin) were cultured in Minimum Essential Medium α (MEMα; Thermo Fisher Scientific). K562 cells (ATCC; female origin) and CSTX002 cells (provided by CYTOSEN; female origin) were cultured in Iscove’s Modified Dulbecco’s Medium (IMDM; Thermo Fisher Scientific), and Raji cells (UC Berkeley Cell Culture Facility; male origin) were cultured in RPMI-1640 (Thermo Fisher Scientific). Unless otherwise specified, media were supplemented with 5% fetal bovine serum (FBS; Omega Scientific) for K562, CSTX002, and Raji cells, or 10% FBS for all other cell lines, as well L-glutamine (0.2 mg/mL; Sigma-Aldrich), penicillin (100 U/L) and streptomycin (100 μg/mL; Thermo Fisher Scientific), gentamycin sulfate (10 μg/mL; Lonza), β-mercaptoethanol (50 μM; EMD Biosciences), and HEPES (20 mM; Thermo Fisher Scientific). Cells were maintained at 37°C in a humidified incubator with 5% CO₂. NK-92 and NK-92 dCas9-KRAB cultures were additionally supplemented with recombinant human IL-2 (100 IU/mL). NK-92 dCas9-KRAB cultures were supplemented with blasticidin (10 μg/mL; InvivoGen) for 1 week of initial culture. All cell lines were routinely tested and confirmed to be negative for mycoplasma contamination.

### CRISPR-mediated editing of mouse NK cells via Cas9 RNP nucleofection

Recombinant Cas9 protein was obtained from the UC Berkeley Macro Lab Core (40 μM Cas9 in 20 mM HEPES-KOH, pH 7.5, 150 mM KCl, 10% glycerol, 1 mM DTT). For *in vitro* functional assays and adoptive transfer experiments, CRISPR guide RNAs (sgRNAs) targeting the murine *Itpkb* gene were designed using the predesigned Alt-R™ CRISPR-Cas9 guide RNA tool (Integrated DNA Technologies, IDT). Corresponding crRNAs were duplexed with tracrRNA (IDT) in duplex buffer at 37°C for 30 min to generate 80 μM sgRNA. For CAR-NK cell experiments, a pool of three sgRNAs targeting murine *Itpkb* was purchased from EditCo Bio and reconstituted in nuclease-free TE buffer to 80 μM. Guide RNA sequences are provided in Table 3.

Splenocytes were isolated and cultured in RPMI-1640 supplemented with 10% FBS, L-glutamine (0.2 mg/mL), penicillin (100 U/mL) and streptomycin (100 μg/mL), gentamycin sulfate (10 μg/mL), β-mercaptoethanol (50 μM), HEPES (20 mM), and recombinant human IL-2 (1000 IU/mL), at a density of 15 mL per mouse. On day 3 of culture, NK cells were enriched using the MojoSort™ Mouse NK Cell Isolation Kit (BioLegend) and subsequently cultured at 2 × 10 cells/mL in the presence of 1000 IU/mL IL-2. On day 5, 5 × 10 NK cells per condition were prepared for nucleofection.

Cas9 ribonucleoprotein (RNP) complexes were assembled at a 1:2 molar ratio by combining 1.25 μL of 40 μM Cas9 with 1.25 μL of 80 μM sgRNA, supplemented with 1 μL of 125 mg/mL poly(L-glutamic acid) sodium salt (Alamanda Polymers). RNP complexes were incubated at 37°C for 15 min and transferred to a single well of a 96-well nucleofection cuvette (Lonza). NK cells were resuspended in 17.5 μL of supplemented P3 buffer from P3 Primary Cell 96-well Nucleofector Kit (Lonza), mixed with 2.5 μL of Cas9 RNP complex, and nucleofected using the CM-137 program on the 4D-Nucleofector (Lonza). Immediately following nucleofection, 80 μL of pre-warmed RPMI-1640 containing 10% FBS was added, and cells were allowed to recover at 37°C for 15 min before being returned to culture in RPMI-1640 supplemented with 10% FBS and 1000 IU/mL IL-2 at a density of 5 × 10 cells/mL.

Edited NK cells were used for functional assays or adoptive transfer experiments 1-3 days after nucleofection. To assess genome editing efficiency, genomic DNA was extracted using QuickExtract™ DNA Extraction Solution (Biosearch Technologies) and used as a template for PCR amplification of the targeted locus. PCR products were subjected to Sanger sequencing at the UC Berkeley DNA Sequencing Facility, and resulting chromatograms were analyzed using TIDE^63^ or ICE^91^ software.

### Flow cytometry

Single-cell suspensions from spleens were prepared by mechanical dissociation followed by passage through a 40-μm cell strainer. Red blood cells were lysed from splenocyte preparations using ammonium-chloride-potassium (ACK) lysis buffer (prepared in-house). Tumors were minced with a razor blade and dissociated using a gentleMACS Dissociator (Miltenyi Biotec), followed by filtration through a 70-μm cell strainer. Tumor-derived single-cell suspensions were subsequently enriched for lymphocytes using Lympholyte-M (Cedarlane) according to the manufacturer’s instructions, washed, and used for downstream flow cytometry assays.

For flow cytometry, single-cell suspensions were incubated for 10-15 min with supernatant from the 2.4G2 hybridoma (produced in-house) to block FcγRII/III, followed by staining with fluorochrome- or biotin-conjugated antibodies for 20 min at 4°C in PBS supplemented with 1% FBS (FACS buffer). Cells that were stained with biotin-conjugated antibodies were subsequently incubated with fluorophore-conjugated streptavidin (BioLegend) for 20 min at 4°C in FACS buffer. For intracellular staining, cells were fixed and permeabilized using the Cytofix/Cytoperm kit (BD Biosciences) and stained with intracellular antibodies for 30 min at 4°C in Perm/Wash buffer (BD Biosciences). For phospho-flow cytometry, cells were fixed by adding an equal volume of prewarmed Cytofix buffer (BD Biosciences) and incubating at 37°C for 10 minutes. Cells were then permeabilized with Perm Buffer III (BD Biosciences) and stored in Perm Buffer III at -20°C until flow cytometric analysis. On the day of analysis, cells were stained with phospho-specific antibodies for 30 min at room temperature in FACS buffer. Data were acquired on an LSR Fortessa, LSR Fortessa X20, or Symphony A3 flow cytometer (BD Biosciences) and analyzed using FlowJo software (Tree Star). Dead cells were excluded using DAPI (BioLegend) or fixable viability dyes (Thermo Fisher Scientific) according to the manufacturer’s instructions. In selected experiments, NK cells were sorted using a FACSAria Fusion cell sorter (BD Biosciences).

### NK responsiveness assay

To assess NK cell responsiveness *ex vivo*, 96-well high-binding flat-bottom plates (Corning) were coated overnight at 4°C with PBS containing the indicated plate-bound antibodies against NK cell activating receptors. Antibodies were used at the following concentrations: anti-NK1.1 (PK136; 50 μg/mL; BioLegend), anti-NKG2D (MI-6; 5 μg/mL; Thermo Fisher Scientific), anti-NKp46 (29A1.4; 5 μg/mL; BioLegend), and anti-NKp30 (MAB18491; 10 μg/mL; R&D Systems). Plates were washed twice with PBS immediately before stimulation. Single-cell suspensions were prepared from the indicated tissues and cultured in antibody-coated wells for 4-5 hours (murine NK cells) or 6 hours (NK-92 cells and primary human NK cells) in the presence of monensin and brefeldin A (both at 1:1000; BioLegend) and fluorophore-conjugated anti-CD107a antibodies (anti-mouse CD107a, 1D4B, 1:500, BioLegend; anti-human CD107a, H4A3, 1:100, BD Pharmingen). Monensin, brefeldin A, and fluorophore-conjugated anti-CD107a antibodies were included at the start of stimulation. Cells were incubated at 37°C in 5% CO₂ during stimulation. For murine NK cell assays, FcγRII/III receptors were blocked by incubating cells for 15 minutes with supernatant from the 2.4G2 hybridoma (produced in-house) prior to plating. In experiments using CRISPR-edited murine NK cells generated by Cas9 RNP nucleofection or splenocytes persistently stimulated with combination of plate-bound antibodies, recombinant human IL-2 (1000 IU/mL) or mouse IL-15 (1.5 ng/mL), respectively, was included during stimulation. Following stimulation, cells were stained with surface markers to identify NK cells, fixed and permeabilized, and subjected to intracellular cytokine staining for IFN-γ. Samples were analyzed by flow cytometry.

### *Ex vivo* cytotoxicity assay

Cytotoxic activity by NK-92 cells and primary human NK cells was assessed using a standard 4-hour ^51^Cr-release assay. Target cells (K562 or Raji) were labeled with ^51^Cr for 1 hour, washed four times with culture medium, and plated at 10^4^ cells per well in 96-well V-bottom plates. Effector NK cells were added at the indicated effector-to-target (E:T) ratios in duplicate or triplicate. Plates were briefly centrifuged prior to incubation to promote effector-target contact and incubated for 4-5 hours at 37°C in cytokine-free medium. Supernatants were then harvested, and ^51^Cr release was measured. Percent specific lysis was calculated as: % specific lysis = 100 × (experimental release - mean spontaneous release) / (mean maximum release - mean spontaneous release), where spontaneous release corresponds to target cells cultured in medium alone and maximum release was obtained by addition of Triton X-100 (final concentration 0.5%).

### Isolation and expansion of NK cells from human peripheral blood

Buffy cones from de-identified healthy adult donors were purchased from Our Blood Institute. Buffy cones were obtained under protocols approved by the supplier. Donor sex was not available. Peripheral blood mononuclear cells (PBMCs) were isolated by density gradient centrifugation using Ficoll-Paque PLUS (GE Healthcare). NK cells were enriched from PBMCs using the EasySep Human NK cell isolation kit (STEMCELL Technologies), according to the manufacturer’s instructions. For NK cell expansion, CSTX002 feeder cells were irradiated twice with 5,000 rad prior to co-culture. Purified NK cells were co-cultured with irradiated CSTX002 cells following an established feeder-based expansion protocol^92^. Briefly, on day 0, NK cells were plated with CSTX002 cells at an E:T ratio of 1:2 at a density of 3.75 x 10^5^ cells / mL in PBMC culture medium consisting of RPMI-1640 supplemented with 10% FBS, L-glutamine (0.2 mg/mL), penicillin (100 U/mL) and streptomycin (100 μg/mL), gentamycin sulfate (10 μg/mL), β- mercaptoethanol (50 μM), and HEPES(20 mM), in the presence of 50 IU / mL IL-2. Complete medium changes were performed on days 3 and 5 post culture. On day 7, remaining cells were counted and replated at 5 x 10^5^ cells / mL in PBMC culture medium supplemented with 500 IU / mL IL-2 for an additional 2-3 days. Efficient elimination of CSTX002 feeder cells and enrichment of NK cells (> 95% CD56^+^CD3^-^) were confirmed by flow cytometry prior to downstream functional assays.

### Generation of lentiviral constructs and lentiviral transduction

To generate a lentiviral vector encoding dual guide RNAs targeting *TRAC* or *ITPKB*, guide sequences were selected by using the CRISPICK platform (Broad Institute) with the human GRCh38 reference genome. Guide RNA sequences are provided in Table 3. Oligonucleotides encoding two guide RNAs per construct were assembled into lentiviral CRISPR interference (CRISPRi) backbone plasmids (pLGR134 and LGR Gibson Dual Guide Insert plasmids; Laboratory for Genomics Research) following BsmBI-v2 digestion (New England Biolabs) and NEBuilder HiFi DNA Assembly (New England Biolabs). For *ITPKB* overexpression and restoration, the open reading frame sequences of human *ITPKB* was cloned into a pHAGE-EF1α-IRES-Thy1.1 lentiviral backbone (modified from a construct provided by Tjian lab, UC Berkeley). The *ITPKB* coding sequence was amplified from PBMC-derived NK cell cDNA and assembled into the vector digested with NotI-HF and BamHI-HF (New England Biolabs) by using NEBuilder HiFi DNA Assembly. An empty vector control was generated in parallel. Lentiviral particles were produced by transfecting HEK293T cells with psPAX2, the lentiviral transfer plasmid, and a BaEV-TR baboon envelope plasmid (provided by E. Verhoygen) using TransIT-LT1 (Mirus Bio). Viral supernatants were collected 48 hours post-transfection, clarified by centrifugation, filtered (0.45 µm), and concentrated by PEG precipitation. Concentrated viral pellets were resuspended in Opti-MEM (Thermo Fisher Scientific). For transduction, viral supernatants were immobilized on 24-well plates coated with retronectin (10 µg/mL; Takara Bio) by spinoculation (1,000g, 32°C, 1 hour). 3 hours later, NK-92 or NK-92 dCas9-KRAB cells (5 x 10^5^ per well) were added in NK-92 medium supplemented with IL-2 (500 IU/ml) and protamine sulfate (20 µg/mL), followed by a second spinoculation (1,000g, 37°C, 1 hour). Media were replaced the following day. For CRISPRi experiments, transduced cells were selected with puromycin (2 µg/mL; InvivoGen) starting 48 hours post transduction. For *ITPKB* overexpression and restoration experiments, Thy1.1-expressing cells were purified by flow cytometry and expanded in IL-2-containing medium prior to downstream functional assays. Knockdown or overexpression efficiency was confirmed by qRT-PCR.

### Desensitization

To induce NK cell desensitization, multiple established models of persistent activation were used, as indicated below.

### Ionomycin-induced desensitization (murine NK cells)

Splenocytes or CRISPR-edited NK cells were treated with 1 µM ionomycin for 22 hours. Cultures were maintained in the presence of IL-2 at 50 IU/mL for splenocytes or 1000 IU/mL for CRISPR-edited NK cells. Following treatment, cells were washed twice with PBS prior to downstream functional assays.

### Persistent stimulation by plate-bound activating receptor ligation (murine NK cells)

Splenocytes were stimulated with plate-bound antibodies targeting NK activating receptors. Anti-NKp46 (29A1.4; 5 μg/ mL; BioLegend) was combined with either anti-NKG2D (MI-6; 5 μg/mL; Thermo Fisher Scientific) or anti-DNAM1 (10E5; 5 μg/mL; BioLegend), coated overnight on 96-well flat-bottom high-binding plates. Cells were plated at 1 x 10^6^ cells per well in 200 μL RPMI-1640 supplemented with 5% FBS, L-glutamine(0.2 mg/mL), penicillin (100 U/ml) and streptomycin (100 μg/mL), gentamycin sulfate (10 μg/mL), β-mercaptoethanol(50 μM), and HEPES (20 mM), in the presence of 1.5ng/ml IL-15. Stimulation was performed for 48 hours.

### Ionomycin-induced desensitization (human NK cells)

NK-92 cells were treated with 1 μM ionomycin for 22 hours at a density of 2.5 x 10^5^ cells/mL in the presence of 200 IU/mL IL-2. Primary human NK cells isolated from PBMCs and expanded on CSTX002 were treated with 1 μM ionomycin for 16 hours at 5 x 10^5^ cells/mL in the presence of 500 IU/mL IL-2. Cells were washed twice with PBS prior to functional assays.

### Desensitization by repeated target cell challenge (CRISPRi-edited NK-92 dCas9-KRAB cells)

To model chronic target cell engagement, NK-92 cells were subjected to repeated co-culture with irradiated K562 cells. K562 cells were irradiated twice at 5,000 rad either on the day before or the day of co-culture. CRISPRi-edited NK-92 dCas9-KRAB cells (2.5 x 10^5^) were co-cultured with irradiated K562 cells at a 1:4 effector-to-target ratio (1 x 10^6^ K562 cells). On days 2 and 4 (or 5), remaining cells were counted and up to 2.5 x 10^5^ cells were re-challenged with freshly irradiated K562 cells at the same ratio. On day 7, cells were again counted and re- challenged at a 1:1 effector-to-target ratio. Co-cultures were maintained at a density of 2.5 x 10^5^ cells/mL in the presence of 100 IU/mL IL-2.

### RNA-seq

For RNA-seq studies, FACS-purified NK cells from each condition were lysed in RLT buffer (Qiagen) supplemented with β-mercaptoethanol and stored at -80°C until processing. 3 biological replicates per condition were analyzed. Total RNA was isolated using the miRNeasy Micro Kit (Qiagen) according to the manufacturer’s protocols. RNA integrity was assessed by using bioanalyzer prior to library preparation. mRNA-seq libraries were generated using the Kapa mRNA HyperPrep kit (Roche). Libraries from TP, SP, and TN CD27^-^CD11b^+^ NK cells isolated from *B2m*^+/+^ or *B2m*^−/−^ mice were sequenced on an Illumina HiSeq 4000 platform using single-end 50bp reads. Libraries for all the other experimental conditions were sequenced on an Illumina NovaSeq X platform using single-end 100bp reads. Sequencing depth ranged from approximately 10-30 million reads per sample. Raw sequencing data were processed using Illumina CASAVA to generate FASTQ files. Reads were aligned to the mouse reference genome GRCm39 by using STAR aligner^93^ with the following parameters: “--outFilterType BySJout --outFilterMultimapNmax 20 --alignSJoverhangMin 8 --alignSJDBoverhangMin 1 --outFilterMismatchNmax 999 --outFilterMismatchNoverReadLmax 0.04 --alignIntronMin 20 -- alignIntronMax 1000000 --alignMatesGapMax 1000000”. Gene- and transcript-level quantification was performed using RSEM^94^. Downstream analyses were performed with R 4.5.2^95^ using Bioconductor 3.22^96^. Genes with low expression were filtered such that only genes with normalized counts ≥5 in at least three samples, or with summed normalized counts across all samples exceeding the number of conditions, were retained for analysis. Differential gene expression analysis was performed using DESeq2^97^. For comparisons involving the CIN and B2m datasets, differentially expressed genes were defined by FDR < 0.1, reflecting increased heterogeneity and limited sample size in these conditions. For all other comparisons, FDR < 0.05 was used as the threshold. Differential expression analyses were performed only between samples generated on the same sequencing platform, with the exception of the comparison between NK cells isolated from spleen vs. tumor. Because spleen- and tumor-derived NK cell samples were sequenced on different platforms, this comparison was used for qualitative validation - specifically to assess directional consistency and enrichment of the previously defined desensitization signature - rather than for unbiased differential gene expression discovery. Gene set enrichment analysis was performed using GSEA^98^ software.

### Antibodies

For flow cytometry, we used the following antibodies.

Mouse: Anti-CD3ε (145-2C11), anti-CD4 (GK1.5), anti-CD8α (53-6.7), anti-CD11b (M1/70), anti-CD19 (6D5), anti-CD27 (LG.3A10), anti-CD45 (30-F11), anti-CD49a (HMα1), anti-CD49b (DX5), anti-F4/80 (BM8), anti–I-A/I-E (M5/114.15.2), anti-Ly6G (1A8), anti-NK1.1 (PK136), anti-NKp46 (29A1.4), anti-Ter119 (TER-119), anti-CD107a (1D4B), anti-IFN-γ (XMG1.2), and anti-Perforin (S16009A) were purchased from BioLegend. Anti-Granzyme A (GzA-3G8.5), anti-Granzyme B (NGZB), and anti-Ly49I (YLI-90) were purchased from Thermo Fisher Scientific. Anti-NKG2A/C/E (20d5) was purchased from BD Biosciences. Ly49C antibody 4LO3311 (a gift from S. Lemieux) was purified and conjugated to biotin according to standard methods. Biotin-conjugated antibodies were detected with streptavidin (BioLegend).

Human: Anti-CD56 (5.1H11), anti-CD16 (3G8), anti-CD3 (OKT3), anti-IFN-γ (4S.B3), and anti-phospho-ERK1/2 (Thr202/Tyr204) were purchased from BioLegend. Anti-CD107a (H4A3) was purchased from BD Biosciences. Anti-phospho-AKT1 (Ser473) was purchased from Thermo Scientific. All antibodies were fluorophore-conjugated unless otherwise specified.

### Ca^2+^ flux assay

NK-92 cells desensitized by treatment with 0.5 μM ionomycin or vehicle-treated control cells were washed twice with Hank’s Balanced Salt Solution (HBSS; Thermo Fisher Scientific) containing calcium chloride and magnesium chloride and supplemented with 1% FBS. Cells (4.5 × 10 per sample) were then loaded with Fluo-4 AM (2 μM; Thermo Fisher Scientific) and Fura Red AM (5 μM; AAT Bioquest) in 225 μL HBSS containing calcium and magnesium, supplemented with 1% FBS and probenecid (5 mM; AAT Bioquest) (referred to as calcium assay buffer), and incubated at 37°C for 40 minutes. During dye loading, cells were gently mixed once at 30 minutes by pipetting. Following dye loading, cells were washed twice with calcium assay buffer lacking dye. Cells were then pelleted and resuspended in 50 μL calcium assay buffer containing anti-NKp30 antibody (10 μg/mL; MAB18491, R&D Systems). Plates were incubated on ice for 20 minutes to allow receptor binding. Cells were washed once by addition of 150 μL calcium assay buffer, resuspended in 200 μL calcium assay buffer, and kept on ice until acquisition. Immediately prior to data acquisition, 100 μL calcium assay buffer was added to each well, and samples were equilibrated in a 37°C water bath for 3 minutes. Samples were acquired for a total duration of 240 seconds. Calcium flux was quantified as the ratio of Fluo-4 to Fura Red fluorescence using derived parameters and kinetic analysis. Dual-dye ratiometric analysis was used to minimize variability due to dye loading and cell size. Median fluorescence ratios were used for trace visualization. Calcium responses were quantified by calculating the area under the curve (AUC), and receptor-induced calcium responses were determined by subtracting the AUC of unstimulated (medium) control samples from that of anti- NKp30-stimulated samples. As a quality-control criterion, samples in which NKp30 stimulation failed to produce a detectable increase in AUC above the matched medium control were excluded from pooled quantification. This predefined criterion was applied uniformly across all editing conditions.

### *In vivo* tumor growth experiments

For subcutaneous tumor implantation, MC38 *B2m*^−/−^ cells, cells were washed and resuspended in PBS (Thermo Fisher Scientific) and 4 x 10^6^ tumor cells in a total volume of 50 μl were injected subcutaneously. For SB28m-EGFRvIII tumors, cells were washed and resuspended in HBSS (Thermo Fisher Scientific), mixed 1:1 with Matrigel (Corning), and 3 x 10^5^ cells in a total volume of 50 μL were injected subcutaneously. Tumor dimensions (length, width, and height) were measured using digital calipers, and tumor volume was calculated using the ellipsoid formula: V = (π/6) × length × width × height. For experiments shown in Figure 6A, mice were intravenously injected with 1 × 10 CRISPR-edited NK cells 4 days prior to tumor inoculation. One day later, mice received an intraperitoneal injection of 10 μg H9-MSA. For experiments shown in Figure 6B, mice were treated with GNF362 (Axon Medchem; 200 μg per dose) or vehicle control (water supplemented with 4% Tween-80 and 5% DMSO by volume) via oral gavage every 3 days, beginning 9 days after tumor inoculation. In selected cohorts, NK cells were depleted by intraperitoneal injection of anti-NK1.1 antibody (clone PK136, Leinco Technologies; 200 μg per dose) on days 7 and 8 post tumor inoculation and weekly thereafter. For experiments shown in Figure 6C, mice were intravenously injected with 1.4 × 10 CRISPR- edited CAR-NK cells 13 days after tumor inoculation. Beginning the following day, mice received intraperitoneal injections of 10 μg H9-MSA every 3 days. To deplete T cells, mice were treated with anti-CD8β (clone 53-5.8, Leinco Technologies; 200 μg) and anti-CD4 (clone GK1.5, Leinco Technologies; 200 μg) antibodies administered intraperitoneally 2 days and 1 day prior to CAR-NK cell transfer and weekly thereafter. In all experiments, prior to treatment initiation, tumor-bearing mice were stratified by tumor volume and randomly assigned to treatment groups such that mean tumor sizes were comparable across groups. Mice were euthanized upon reaching institutional humane endpoints.

### RNA isolation, reverse transcription, and quantitative polymerase chain reaction

Total RNA was isolated using the TRIzol reagent (Thermo Fisher Scientific) following the manufacturer’s instructions. Complementary DNA (cDNA) was generated using the HiScript III RT SuperMix for qPCR with gDNA wiper (Vazyme). Quantitative real-time polymerase chain reaction (qRT-PCR) was performed using Taq Pro Universal SYBR qPCR Master Mix (Vazyme) in a CFX384 Thermocycler (Bio-Rad). *GAPDH* was used as a reference for human NK cells and *Polr2a* and *Tbp* were used as references for murine NK cells.

Primer sequences were as follows: human *GAPDH*, GGAGCGAGATCCCTCCAAAAT (forward) and GGCTGTTGTCATACTTCTCATGG (reverse); human *ITPKB*, TCTCCTCATCCTACGAAGACTCA (forward) and GCTCACTCTAGGTTTCTGCTGG (reverse); mouse *Polr2a*, CTAAGGGGCAGCCAAAGAAAC (forward) and CCATTCAGCATACAACTCTAGGC (reverse); mouse *Rpl13a*, CTGTGAAGGCATCAACATTTCTG (forward) and GACCACCATCCGCTTTTTCTT (reverse); mouse *Itpkb*, ACTTCCGTGAGAATGGAAAGAGG (forward) and TGAGGGCAGAATCATGTCCAG (reverse).

### Luciferase Assay

25,000 cells / well were lysed in 20 μL passive lysis buffer (Promega) for 15 min at room temperature with gentle rocking on an orbital shaker. The cell lysates were then added with 100 μL Luciferase Assay Reagent (Promega), and luminescence was measured on a SpectraMax L Luminescence Microplate Reader (Molecular Devices). The relative luminescence was calculated as the difference in luminescence between NKp30-stimulated wells and medium controls.

## Author Contributions

Conceptualization: D.H.R. and Y.J. Methodology: Y.J., A.K., S.W.L., B.B.T., and Y.S. Investigation and experiments: Y.J., A.P., Z.G., M.M., E.A., H.S., and C.Z. Writing – original draft: Y.J. and D.H.R. Funding acquisition: D.H.R. Provided critical reagents: H.O., Z.Z., R.A.S., A.N.M., J.D.

## Resource Availability Lead Contact

Requests for further information and resources should be directed to and will be fulfilled by the lead contact, David H. Raulet.

## Materials availability

All unique stable cell lines generated in this study are available from the Lead Contact upon reasonable request.

## Data and code availability

RNA-seq data generated in this study have been deposited in the Gene Expression Omnibus (GEO) under accession number GSE336796 and are publicly available. No custom code was generated in this study. All software used for data analysis is listed in the Materials and Methods. Any additional information required to reanalyze the data reported in this paper is available from the Lead Contact upon request.

## Acknowledgements

We thank L. Zhang and E. Seidel for laboratory support, M. Dupage and T. Mann for providing comments on the manuscript, and the members of the Raulet and Dupage laboratories as well as the Division of Immunology and Molecular Medicine at UC Berkeley for helpful discussions. We thank K. Heydari, M. Delcroix, and H. Dhaliwal of the UC Berkeley Cancer Research Laboratory Flow Cytometry and QB3 Genomics at UC Berkeley (RRID: SCR_022170) for assistance with quality control and quantification of RNA-seq libraries as well as sequencing runs. Part of this work involving HiSeq 4000 platform was support by NIH S10 OD018174 Instrumentation Grant for QB3 Genomics. The plasmid to express dual guide RNAs and NK-92 cells expressing dCas9-KRAB machinery were generously provided by the Laboratory for Genomics Research established by GSK, UCSF, and UC Berkeley. CSTX002 cell line was kindly provided by CYTOSEN.

