## Supplementary Figures for "ITPKB is a conserved regulator of natural killer cell desensitization/education that constrains antitumor immunity"

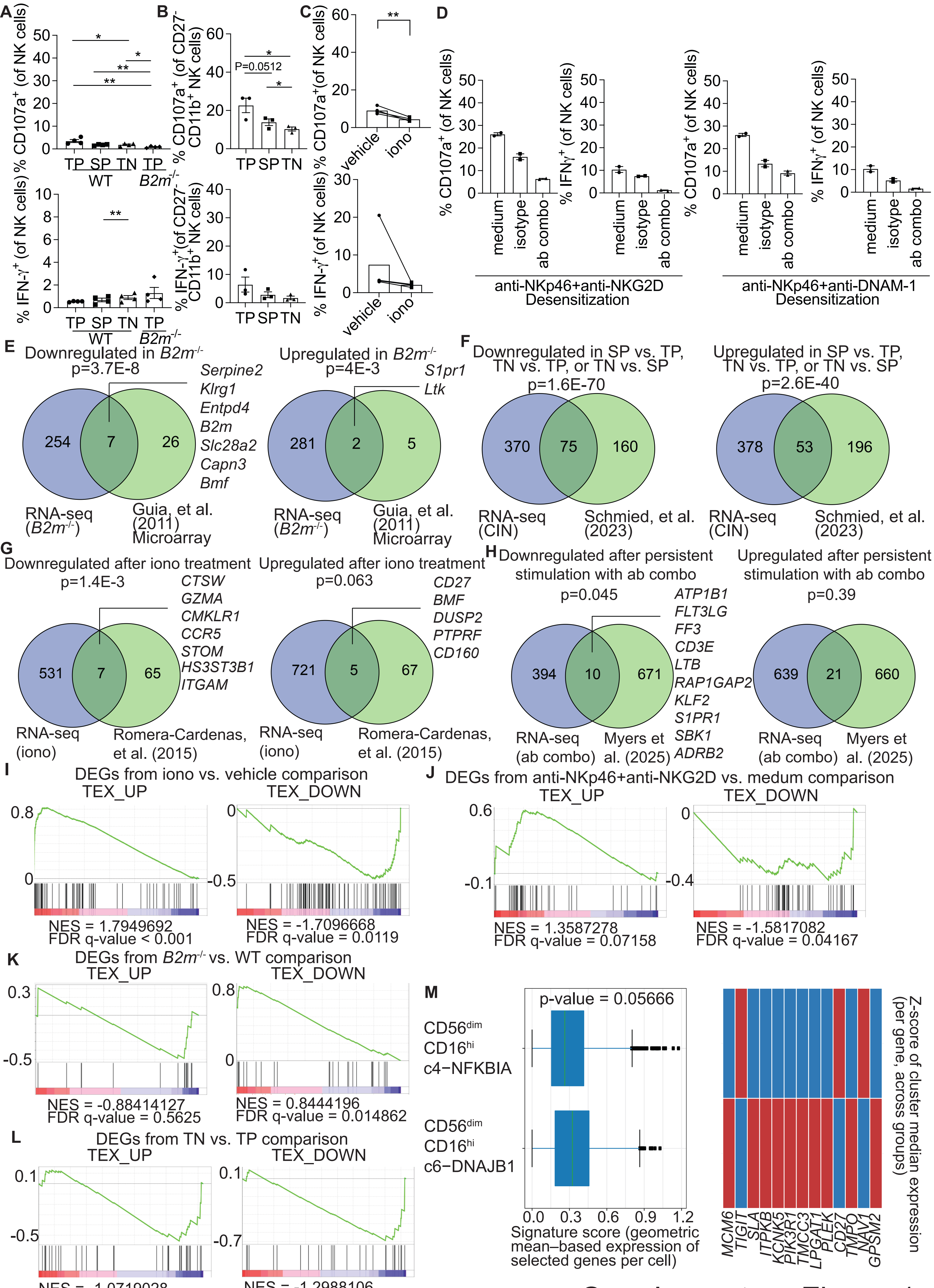


Supplementary Figure 1

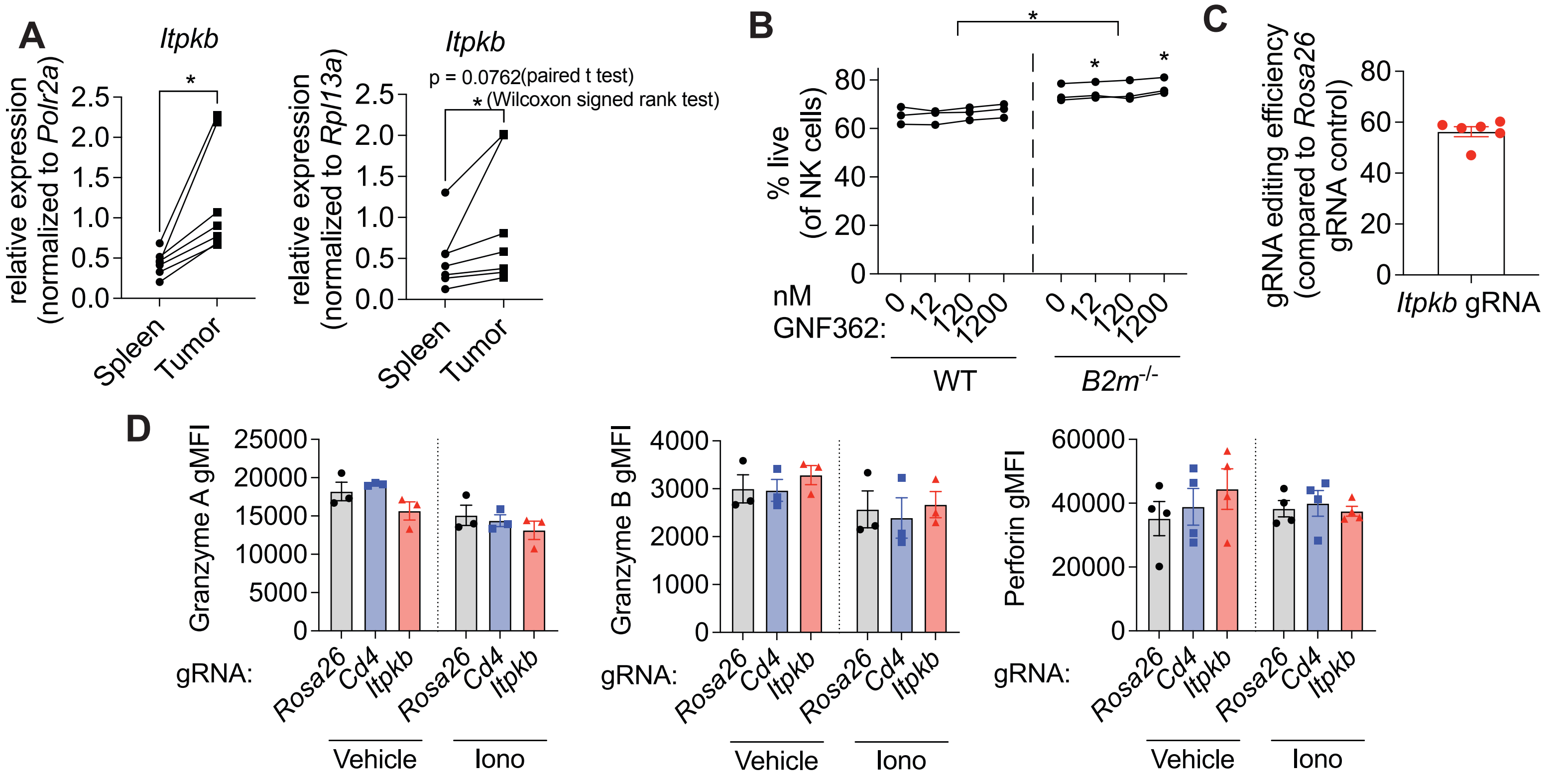


Supplementary Figure 2

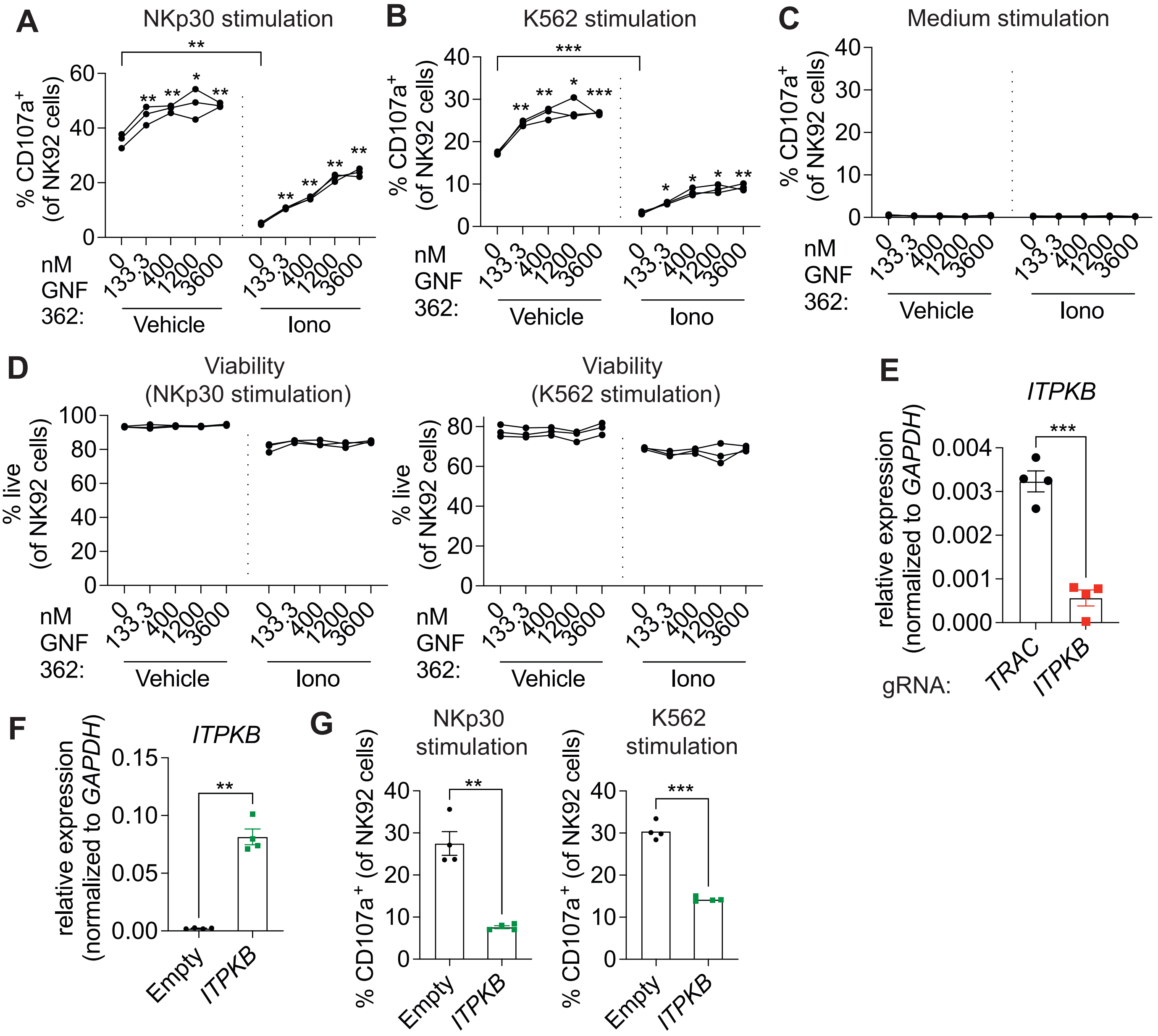


Supplementary Figure 3

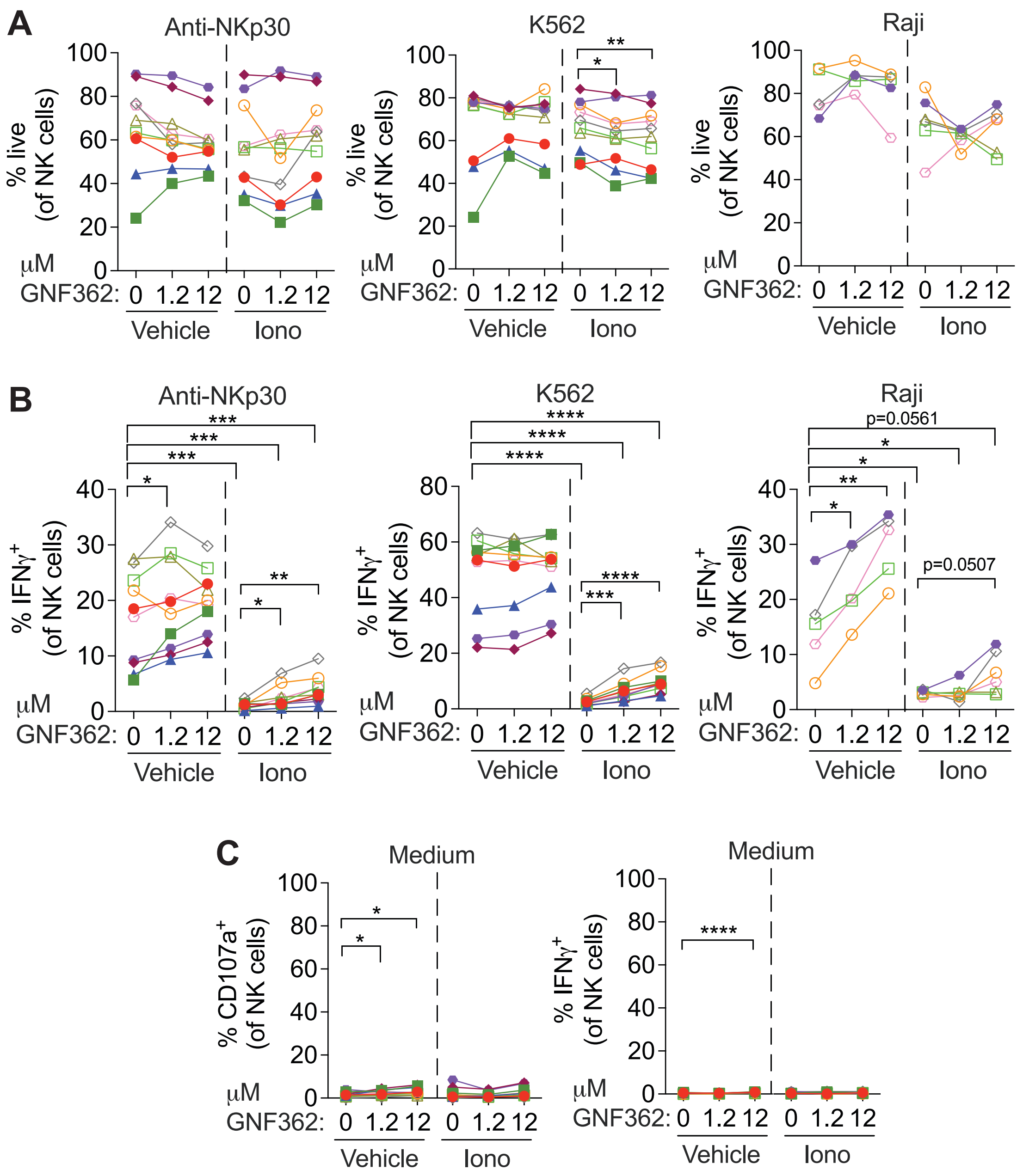


Supplementary Figure 4

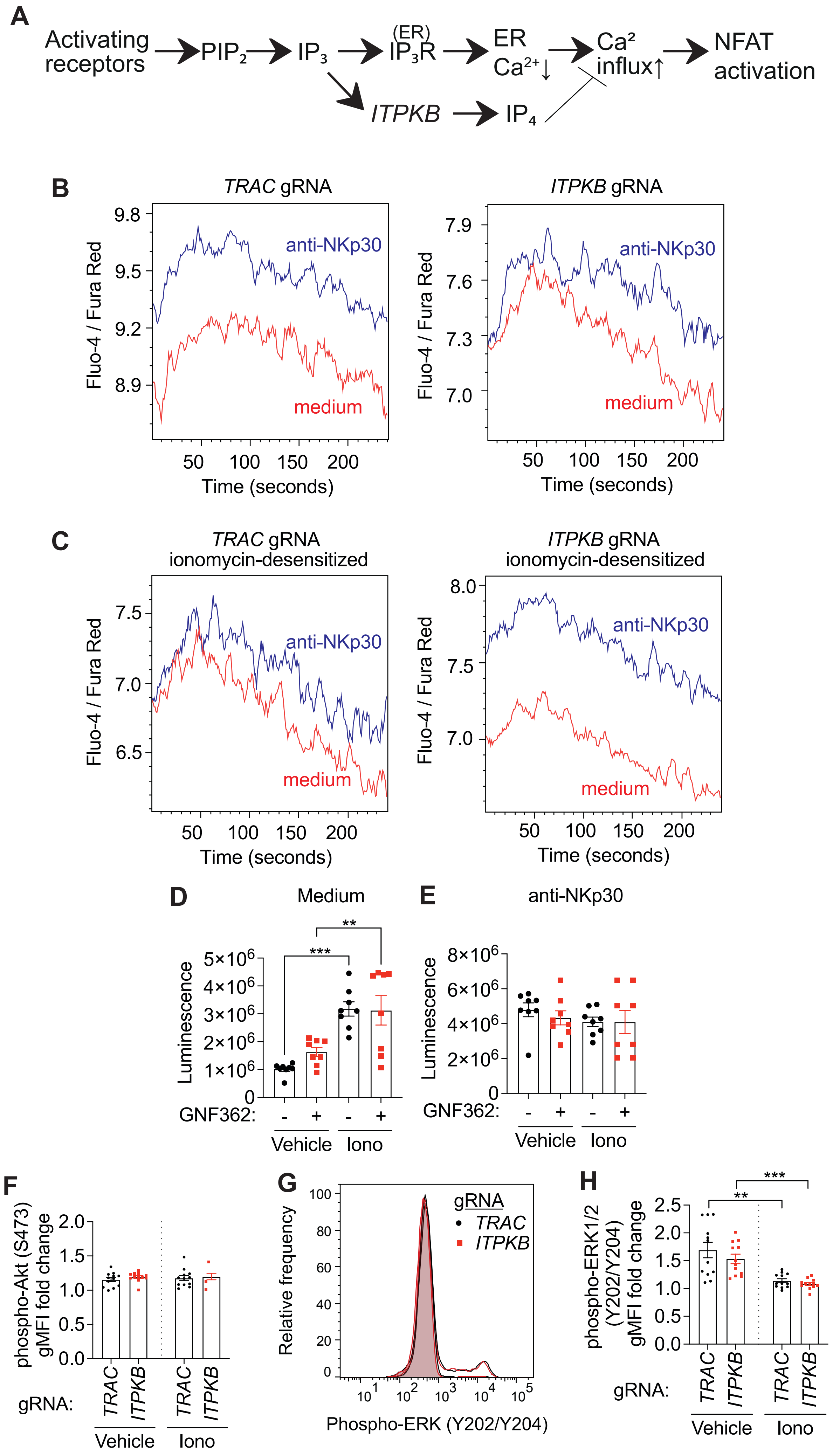


### Supplementary Figure 5

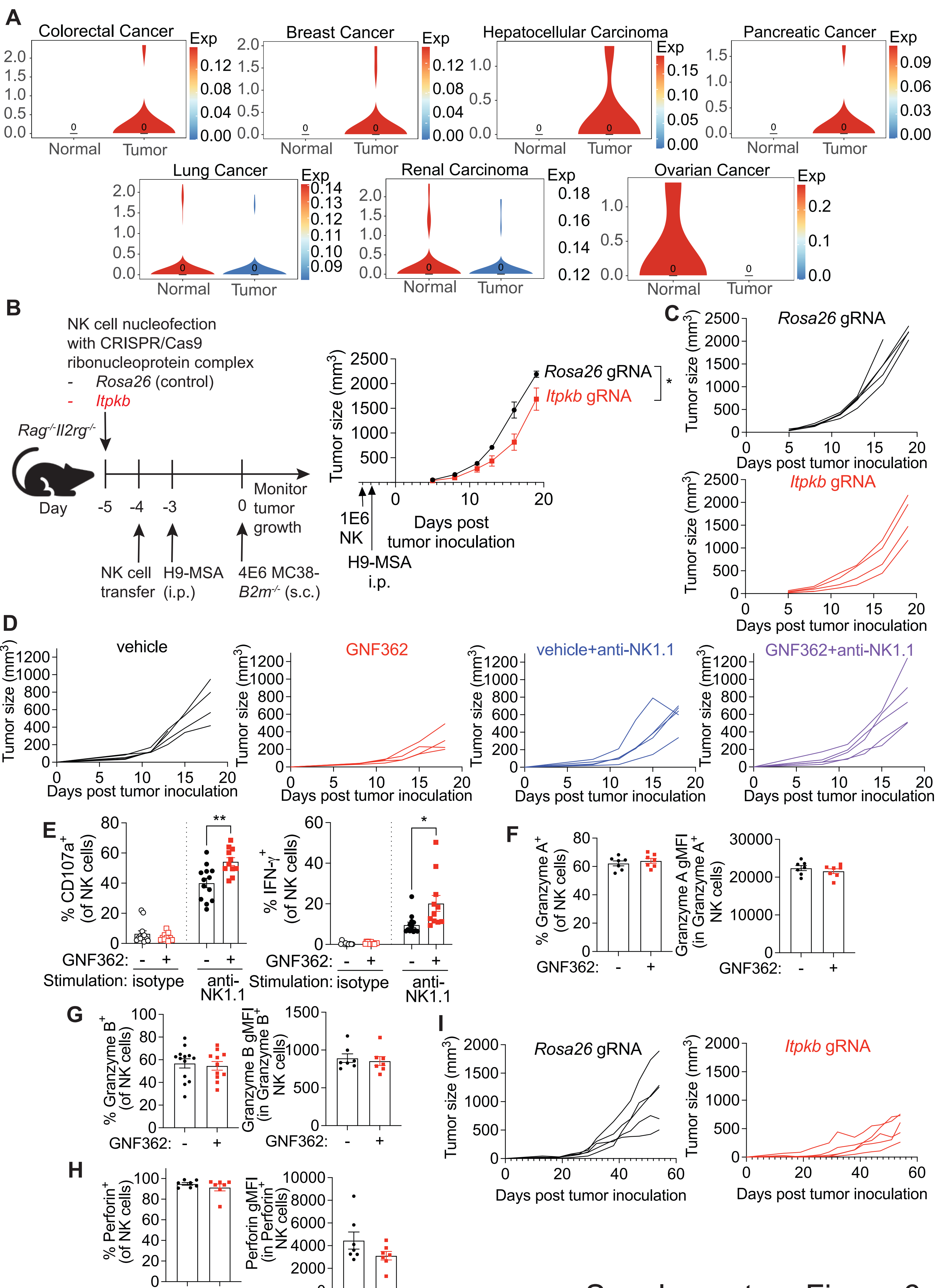


Supplementary Figure 6
